# EMBEDR: Distinguishing Signal from Noise in Single-Cell Omics Data

**DOI:** 10.1101/2020.11.18.389031

**Authors:** Eric Johnson, William Kath, Madhav Mani

## Abstract

While single-cell “omics” based measurements hold the promise of unparalleled biological insight they remain a challenge to analyze owing to their high-dimensional nature. As a result, Dimensionality Reduction (DR) algorithms are necessary for data visualization and for downstream quantitative analysis. The lack of a principled methodology for separating signal from noise in DR algorithmic outputs has limited the confident application of these methods in unsupervised analyses of single-cell data, greatly hampering researchers’ ability to make data-driven discoveries. In this work we present an approach to quality assessment, **EMBEDR**, that works in conjunction with any DR algorithm to distinguish signal from noise in dimensionally-reduced representations of high-dimensional data. We apply EMBEDR to t-SNE- and UMAP-generated representations of published scRNA-seq data, revealing where lower-dimensional representations of the data are faithful renditions of biological signal in the data, and where they are more consistent with noise. EMBEDR produces easily interpreted p-values for each cell in a data set, facilitating the comparison of different DR methods and allowing optimization of their global hyperparameters. Most compellingly, EMBEDR allows for the analysis of single-cell data at a single-cell resolution, allowing DR methods to be used in a cell-wise optimal manner. Applying this technique to real data results in a biologically interpretable view of the data with no user supervision. We demonstrate the utility of EMBEDR in the context of several data sets and DR algorithms, illustrating its robustness and flexibility as well as its potential for making rigorous, quantitative analyses of single-cell omics data. EMBEDR is available as a Python package for immediate use.

## Introduction

Advances in high-throughput measurement techniques are revolutionizing biology. The advent of single-cell omics approaches, in particular, promises to illuminate the processes of cellular differentiation, multicellular patterning, signaling, and variation at single-cell resolution [1–13]. However, omics data is high-dimensional — each measured gene adds a dimension to the sample space — leading to an explosive increase in the volume occupied by the data due to the curse of dimensionality (See Figure S1 for an illustration) [14]. In addition, single-cell methodologies generate significant noise due to the small amount of material being measured [15–19]. Thus, despite the great promise that single-cell omics approaches hold it remains a challenge [20] to separate signal from noise in these data sets or make data-driven inferences [14].

Faced with the challenges posed by high-dimensional data sets, a host of sophisticated methods have been developed to help make quantitative inferences from the data. One such class of methods, termed **dimensionality reduction** (DR), attempts to reduce the size (dimensionality) of the data by identifying a reduced set or combination of features (genes) on which further qualitative or quantitative analysis can be applied with more power. Significant effort has been put into the development and application of DR algorithms such as PCA [21], t-SNE [22], UMAP [23], and others [24–38]. Each of these methods attempts to find a lower-dimensional (usually two- or three-dimensional) representation, or *embedding,* of the data that preserves important aspects of the original data structure (for a review, see [39–41]; in application to omics data, see [42]).

Ideally, a researcher would like to use a dimensionally-reduced representation of the data to gain biological insight. For example, if cells from a tissue are sequenced, to what extent can we say that two clusters in the embedding correspond to distinct, differentiated, cell-types? If clusters in such a view are connected by a bridge of cells, does this imply the existence of a path along which cells are differentiating? If cells subjected to different treatments of a drug are processed through a DR method, how is the strength of the treatment effect correlated with distance in the lower-dimensional space? Experimentally, one might be concerned with the depth of sequencing or the number of samples; how does this information get transformed into a dimensionally-reduced representation of the data? Put plainly, DR methods produce an approximate picture of the data, and we’d like to know what parts display biological signal, and what parts are simply noise.

In traditional data analyses, statistics provides a rigorous framework with which to answer these questions, but DR methods confound the statistical distinction between signal and noise. Specifically, DR methods: generically produce distortions in their representations of data, and these distortions are inhomogeneous across a representation [30, 40, 43–47]; are often stochastic and non-linear, meaning that the robustness and reproducibility of results is hard to assess [41]; and often require user specification of hyperparameters, where this specification is often based on heuristics rather than quantitative principles [10, 48–50]. Addressing these issues provides the motivation for this work, as recovering the ability to separate signal and noise in DR output is essential for their utilization in quantitative analyses.

These difficulties with DR methods can be insidious. As an illustration, consider a sample data set that populates the tips (vertices) of a regular tetrahedron in three dimensions. (A slightly more complicated example can be found in Figure S2.) The vertices of this tetrahedron are all equidistant in the original three dimensions of the data, but any squashing of the pyramid into two dimensions will necessarily result in the distances between some pairs of vertices being distorted. For example, flattening the pyramid onto its base will make the top vertex look artificially close to the other three. Alternatively, moving the top to a point outside the bottom triangle will make it artificially far from one of the base vertices. Real data are more complicated than a tetrahedron: cells are arranged in gene expression space in unknown geometric relationships with heterogeneous densities. But if in even simple cases one cannot match nearby regions in the original data to nearby in the DR output — or far as far — any interpretation of the dimensionally-reduced representations of real single-cell data must proceed with caution.

To address the distortive effect of reducing dimensions, DR algorithms often employ stochastic or non-linear techniques, which work with remarkable success in a variety of contexts [41]. Using these techniques, however, also means that the exact outputs of a DR method will rarely look similar, whether comparing across methods, different parameter choices with the same method, or even across separate runs of the same method with identical choices of parameters. As an example, consider Figure 1, where scRNA-seq data from over 5000 bone-marrow cells from the Tabula Muris Cell Atlas [8] have been embedded in 2D using t-SNE [51] and UMAP [23] each at two different user-prescribed settings. (Throughout this work, we use *k*_Eff_ to parameterize t-SNE instead of `perplexity`. See Section S3 for more information.) In each panel of this figure, the lower-dimensional representations demonstrate some apparently clustered structures, but the number, size, and shape of these clusters varies dramatically between the representations. As an example, the precursor cells in groups 1, 5, and 9 (purple, yellow, and green) may seem to occupy three to four distinct “clusters” in the top panels, but are clearly part of a single cluster in the lower panels. Furthermore, the properties of these putative structures also changes in separate runs of the algorithms, as shown in Figure S4. Without more information then, it is not obvious which of these panels best represents the high-dimensional structure of the data. An assessment of the “error” in these representations would allow for such a determination.

**Figure 1:**
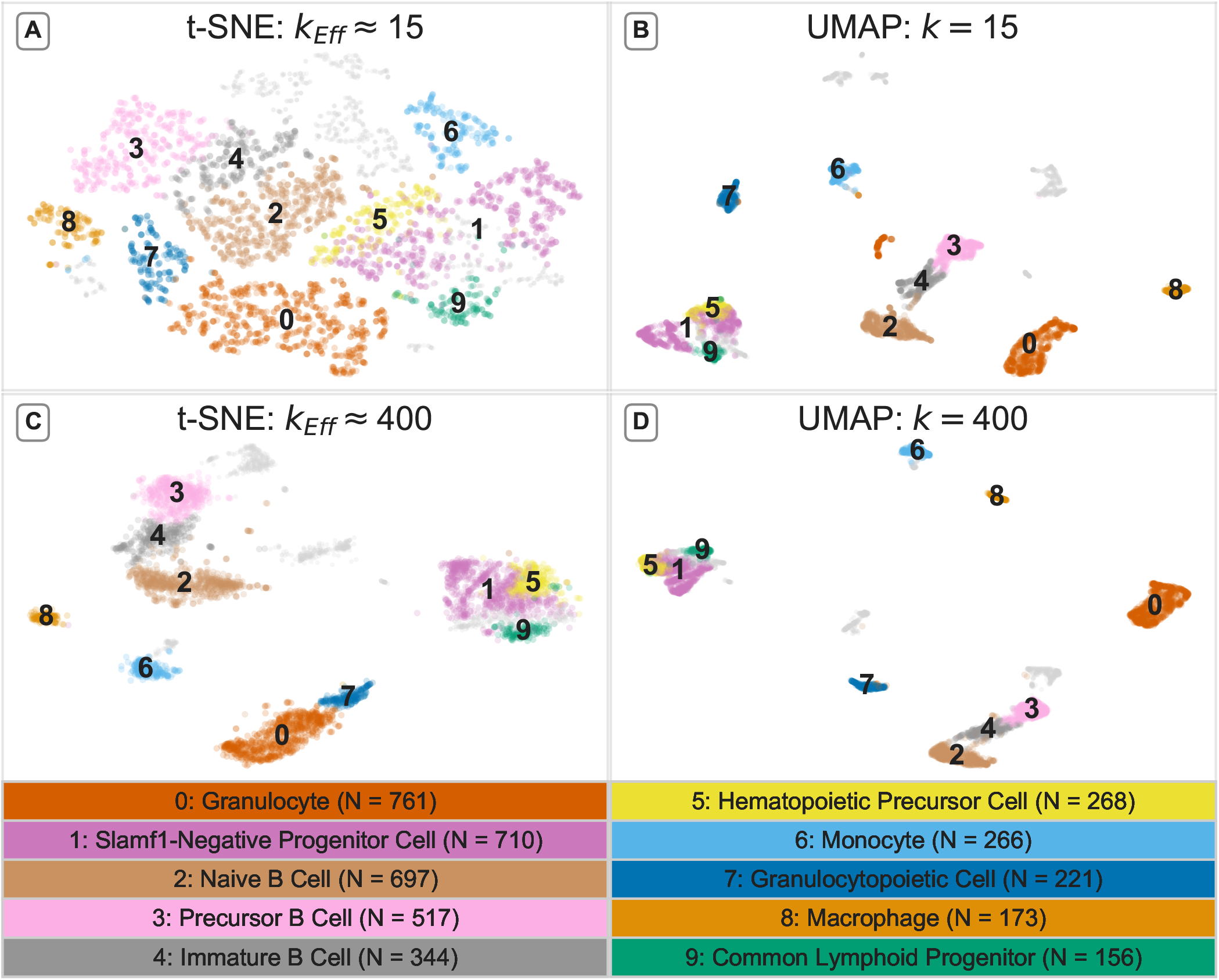
Features of dimensionally-reduced data are sensitive to the choice of algorithm and algorithmic settings: Four dimensionally-reduced representations of RNA-seq measurements from 5,037 bone-marrow cells collected by the Tabula Muris Consortium [8] generated by t-SNE at *k*_Eff_ = 15 (A) and 400 (C) (`perplexity` = 7 and 250, respectively; openTSNE implementation [51]) and by UMAP [23] at `n_neighbors` = *k* = 15 and 400. Ten annotated cell types provided by [8] are colored and labelled. The same cells are colored and labelled in each panel. In (B), the number of nearest neighbors, *k* is set to its default value, 15, in UMAP. For comparison we use t-SNE with the same number of effective nearest neighbors (*k*_Eff_ = 15) in (A), as described in S3. Following the method in S3, we use t-SNE with a similar number of nearest neighbors (*k*_Eff_ = 15) in (A). In panels (C) and (D) we visualize the data using t-SNE and UMAP, respectively, at a much larger number of nearest neighbors: *k*_Eff_ ≈ 400 in (C) and *k* = 400 in (D).

Together, these observations strongly motivate the need for methods to assess the size of DR-induced “error” associated with representing high-dimensional data in lower-dimensional spaces. We emphasize that even noiseless data will be distorted during the DR process, making error assessment a necessary component in applying these methods. More specifically, we assert that a successful error-quantification scheme should achieve the following:

1. **Assess Quality Locally:** Since the errors incurred in reducing the dimensionality of data are not distributed homogeneously across the lower-dimensional representation [45, 46], a quality assessment scheme should provide *local* (per-cell) error estimates as opposed to a single global estimate.
2. **Assess Variability in Quality:** To account for changes in quality that may be due to variation across different executions of a stochastic DR algorithm, a quality assessment scheme should consider the *distribution* of errors across runs.
3. **Assess Quality Statistically:** A robust quality assessment scheme should employ a null hypothesis in order to establish a “ruler” or baseline against which errors in data can be compared.

Others have addressed the problem of DR quality assessment: work has been done to provide heuristic guidelines on how to appropriately use DR algorithms [10, 48–50] and to make improvements to the algorithms themselves [51–58]. Several efforts to characterize the quality of DR methods have been pursued [41, 46, 59], which can roughly be categorized as being global [30, 53, 60–68] or local [29, 45, 46, 69, 70] in scope, and either based on preserving distances [68], neighborhoods [30, 46, 59–61, 63, 71, 72], or topology [67, 73, 74], but in all cases, they attempt to summarize the extent to which a given DR algorithm preserves some aspect of the original data’s structure. In surveying this literature, and considering our basic principles, we find that what is still missing is an approach that not only assesses quality quantitatively and locally [45, 47, 50, 70–73, 75], but also *statistically* in that it seeks to characterize the part of the natural and expected variability in quality that is due to noise.

It is with this in mind that we have developed the **E**mpirical **M**arginal resampling **B**etter **E**valuates **D**imensionality **R**eduction, or **EMBEDR**, algorithm in order to locally and statistically evaluate DR error. EMBEDR is a general approach that addresses the several unique concerns that arise with high-dimensional, noisy data like scRNA-seq measurements, while also adhering to our motivating principles for a quality assessment scheme.

## The EMBEDR Algorithm

In this section we describe the heuristic structure of the EMBEDR algorithm, as well as specific implementation details that are reflected in the figures throughout this work. Considered generally, EMBEDR is based on measuring the local, per cell, distortion of the DR method as a “quality” statistic. We then use empirical resampling methods to generate a null distribution for these statistics so that we may quantitatively assess whether a dimensionally reduced view of a cell’s local neighborhood has more structure (signal) than we expect from noise. We re-emphasize that the EMBEDR framework is agnostic to the DR method being employed and the ways in which quality is assessed. That is, EMBEDR is not designed specifically to evaluate the accuracy of t-SNE and UMAP for sc-omics data, but more generally to assess the quality of any DR method applied to high dimensional data.

The EMBEDR algorithm consists of three steps: **(1)** the repeated *embedding* of the data (the repeated generation of lower-dimensional representations of the data), **(2)** the construction and embedding of null data sets generated in a data-driven manner, and **(3)** the calculation of the quality statistics and performance of a hypothesis test. These are illustrated in Figure 2(A), (B), and (C), respectively. We elaborate on each of these three steps below. As suggested by the motivating principles, these steps focus on the calculation of a local quality statistic, the Empirical Embedding Statistic *(EES*), for each sample (cell) in the data set. We then go on to describe how our algorithm characterizes the distribution of the EES in a meaningful and useful way.

**Figure 2:**
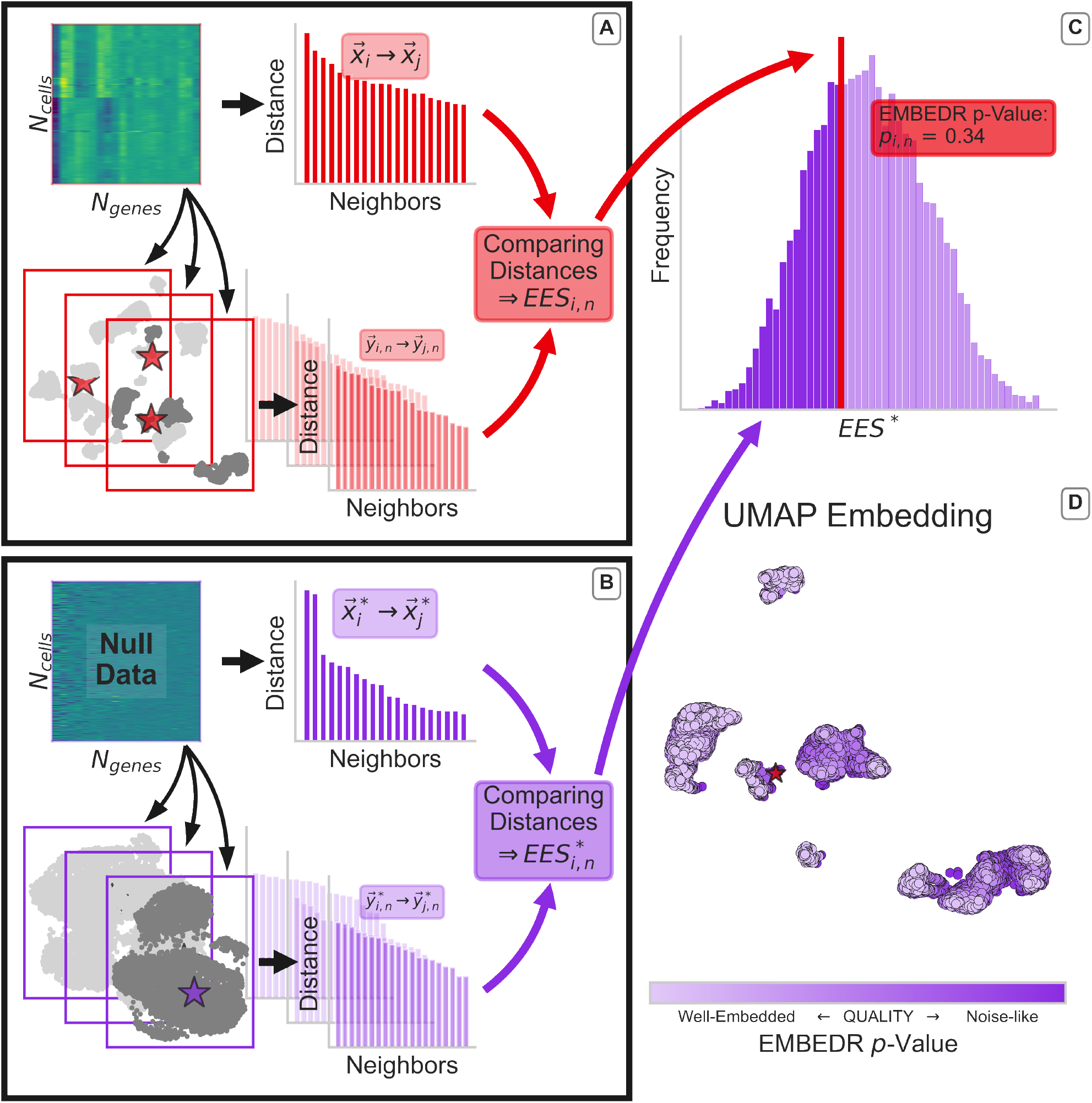
A schematic of the EMBEDR Algorithm: In (A), the data (5037 FACS-sorted marrow cells from [8], shown as a heatmap) are embedded in 2D using a DR method several times (here: UMAP with *k*=n_neighbors=100). For each sample, the distances to neighboring samples are calculated in both the original data, 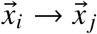, and the low-dimensional embedding, 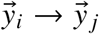. An example cell is illustrated by a red star in the embeddings. These distance distributions are compared to calculate *EES_i,n_*, a quality score for each cell in each embedding. In (B), the same procedure as in (A) is conducted using null data sets constructed via marginal resampling (See Figure 3). In (C), the individual *EES_i,n_* are compared to the null distribution of *EES* * to estimate a *p*-value for each cell’s embedding quality. This *p*-value corresponds to the likelihood that the null data could generate an observed or better embedding quality. In (D), these *p*-values are illustrated as a color, so that embedding quality can easily be visualized across an embedding.

To clarify the notation throughout the rest of this paper: consider a data matrix *X* to be a collection of *N*_cells_ vectors, where each cell contains measurements for each of *N*_genes_ genes. Noting that for stochastic DR algorithms, the data can be embedded multiple times to yield different lower-dimensional representations, we denote the position of the *i*^th^ cell in the *n*^th^ embedding by 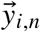, where the number of embeddings is *N*_embed_, and 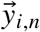 is usually a 2- or 3-D vector. For each cell, in each embedding, we will calculate the quality statistic, which we denote EES_*i,n*_. An asterisk, *, is used to indicate quantities that correspond to “null data” generated by resampling, so that a resampled high-dimensional data vector is 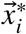 and its position in the embedded space would be 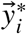. The final step of the hypothesis test process involves calculating the *p*-value: *P_i,n_* =< Prob (EES* ≤ EES_*i,n*_) using an empirically generated EES* distribution. (EES* refers to the set of 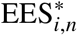 across all cells in the null data and all 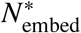 embeddings of the null data.)

1. **Embedding the Data:** The first part of the EMBEDR algorithm is to use a candidate DR algorithm to embed high-dimensional data in lower-dimensions. For stochastic algorithms, such as t-SNE or UMAP, this embedding must be performed multiple times as the quality of a specific sample’s location can vary dramatically between embeddings (see Figure S10). This is illustrated in Figure 2(A) using UMAP. In the final step of the algorithm, the effect of these multiple embeddings will be summarized into a single quantity, so that the choice of *N*_embed_ is not critical to the interpretation of the output (see Supplemental Section S4). Next, an affinity between pairs of cells in the high-dimensional space is calculated by applying a Gaussian kernel with fixed entropy to the pairwise distances (as in [22]). This is repeated in the lower-dimensional embedding except that a Student’s *t*-distribution is used to calculate affinities. The affinity distributions for each cell in high and low dimensions are compared using the Kullback-Leibler Divergence, *D_KL_* [76], which constitutes our quality measurement. If the *D_KL_* is small, it indicates that the two distributions are similar, suggesting that the neighborhood of the embedded cell looks similar to its neighborhood in the original, high-dimensional, gene expression space. This calculation is illustrated in Figure 2(A). The use of *D_KL_* as a quality metric has also been used in other contexts [30, 77]. For more details on how this is calculated, see Section S1.
2. **Null Construction and Embedding:** The most crucial step in the EMBEDR algorithm is the data-driven construction of biologically-realistic “null” data sets that can be used to generate an expectation for embedding quality from data devoid of biological signal. EMBEDR achieves this via *marginal resampling*, which is a resampling procedure where each gene’s expression levels in the null data are independently drawn from the distribution for that gene in the original data. Figure 3 illustrates this process. Computationally, if *X* is an *N*_cells_ × *N_genes_* data matrix of gene expression observations, *X*\* can be generated by independently drawing *N*_cells_ samples from each column in X with replacement (the resulting *X*\* has the same shape as *X*). In this way, the null data contains biologically realistic, marginal, distributions of individual genes — Figure 3(B) shows that genes have nearly *identical* marginal distributions in both data sets — but the joint distribution of genes is altered. More technically, the null data set comprises a joint probability distribution constructed from the explicit product of the individual marginal distributions — guaranteeing independence of the genes in the null data. This property of independence generates a more diffuse distribution of cells relative to the real data, allowing for the assessment of whether real cells populate higher density regions in expression space than expected. Any clustering that manifests in the null data set is a consequence of the properties of the marginal distributions and the algorithm employed. The use of marginal resampling has been used successfully in several other contexts where the signal under examination was assumed to be a result of correlations in the data [78–80]. This is a reasonable assumption for signals that are discoverable by DR methods, as these methods leverage the covariance (PCA) or pair-wise distance (t-SNE, UMAP) matrices to generate embeddings. Constructing null data via marginal resampling is also a model-free and a parameter-free process. In the context of scRNA-seq data, these resampled data sets correspond to the hypothesis that all cells are sampling a common distribution of gene expression, which is a useful and generic null hypothesis for many biologically interesting problems, such as cell-type identification, where the hypothesis would be that gene distributions depend on cell identity. Figure 3(C-D) serves to underscore why we should generate these null data empirically: uncorrelated data are not necessarily uniform, meaning that clusters and structures can appear in DR representations of signal-less data! This is not necessarily intuitive, as one might naïvely expect clustering to be a consequence of cells having similar expression profiles, but clusters will be generated by many DR methods even when no such signal is present. Furthermore, there are no theoretical results that describe the application of arbitrary DR methods to arbitrary data, so that marginal resampling is also a practical approach.
3. **Empirical Hypothesis Test:** The final step in the EMBEDR framework is to perform an empirical hypothesis test. Once the null data have been created and the null embedding statistics *EES*\* have been calculated for every sample over several embeddings, each of the sample statistics, *EES_i,n_*, can be compared to the aggregated distribution of null statistics, as illustrated for a sample point in Figure 2(C). The fraction of null statistics, *EES*\*, that are smaller than *EES_i,n_* can be used to estimate the likelihood that null data would be embedded as well by uncorrelated data. This likelihood is interpreted as an empirical *p*-value, and can be summarized across the *N*_embed_ embeddings [81–83] to give a single quality metric, *pi*, for each cell. For the sake of interpretability, we use the *N*_embed_ embeddings of the data to make an estimate of the likelihood that a cell’s quality is better than that of the null, *P*(*EES_i_* ≤ *EES*\*), which amounts to averaging the individual embeddings’ *p*-values. See Section S4 for more details. The EMBEDR *p*-values can then be used, as in Figure 2(D), to color each cell within an embedding indicating regions of higher or lower amounts of embedding quality. When using *D_KL_* as the quality statistic, lower *p*-values indicate that a cell’s neighbors are similarly distanced in the original and low-dimensional spaces, with closer samples (in the original space) weighted more than those further away. The use of other quality metrics would require an appropriate adjustment to this interpretation, but the interpretation of the *p*-value as a measure of better or worse than noise does not.

**Figure 3:**
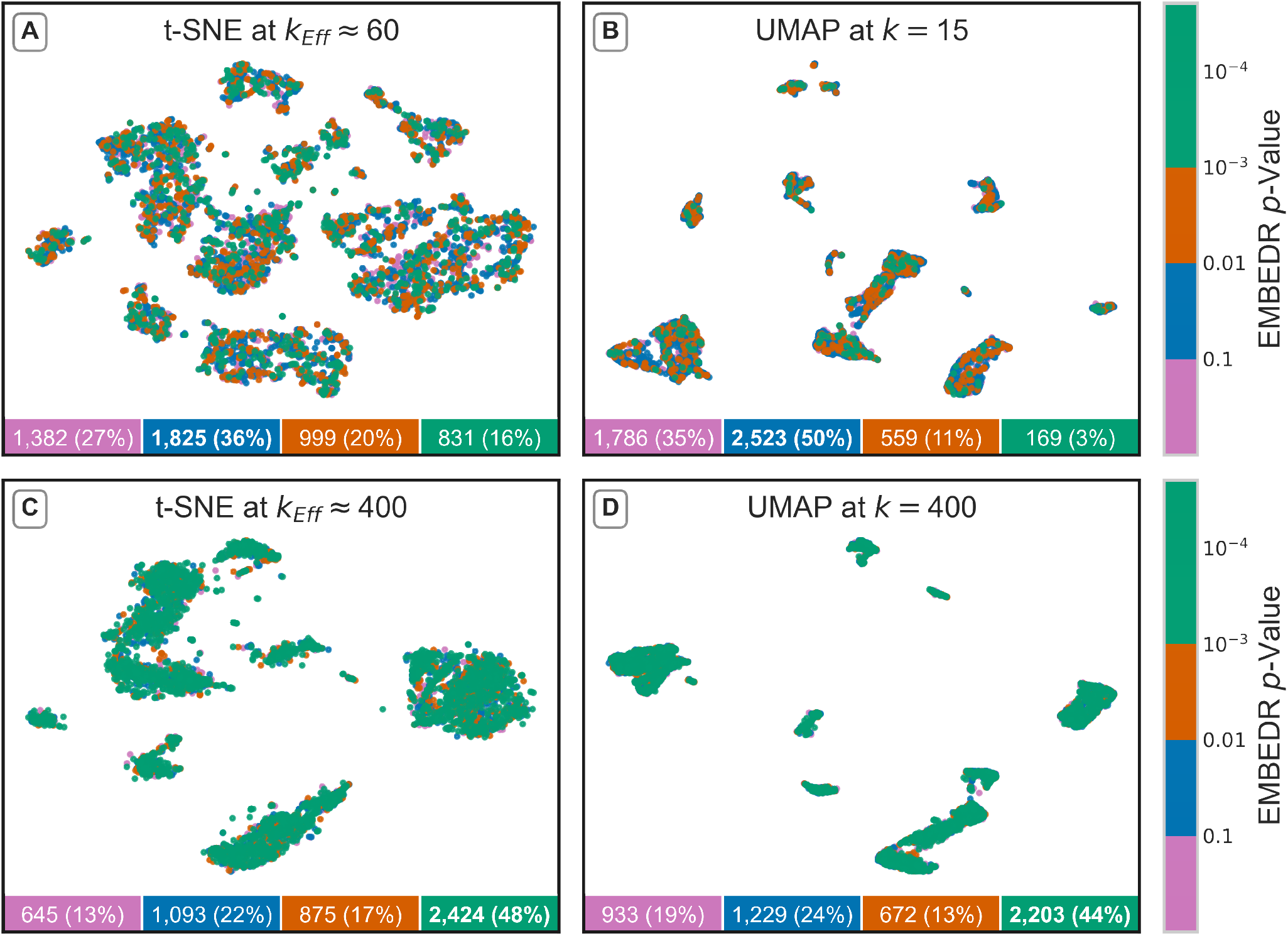
An Overview of Marginal Resampling for Generating Null Data Sets. (A) Gene expression data for real and resampled scRNA-seq data (5037 FACS-sorted marrow cells [8]) are shown as heatmaps. (B) The first and second principal component of the data in (A) are plotted against each other and the corresponding marginal distributions are shown to the top and right. Kernel density estimates (KDEs) are also plotted on the marginal distributions. The effect of marginal resampling to generate null distributions is shown in (C) and (D), where the data and a null data set are embedded using UMAP at *k* = 15 and t-SNE at *k*_Eff_≈ 60, respectively.

In practice, the EMBEDR algorithm operates in conjunction with, not as a substitute for, any DR algorithm, requiring little user input beyond what the DR method would require on its own. The algorithm has been implemented as a ready-to-use Python package on Github for t-SNE, UMAP, and PCA.

## Results

### 1 EMBEDR Reveals Where DR Output Shows Signal vs. Noise

Now that we have a local and statistical approach to separating signal and noise in DR output, we can start to address the difficulties introduced by DR methods in a principled way. For example, we used the tetrahedron thought experiment (Figure S2) to intuitively show how DR methods introduce heterogeneous distortions in the dimensionally-reduced embeddings, but the problem here is not that these methods generate such errors, it’s that they are not systematic or predictable. That is, if there were any system or pattern to misrepresentations in the lower-dimensional embedding, then any of its features, such as the relative separation of two clusters or a cell’s similarity to its neighbors, could be inferred by taking into account those patterns. Of course, single-cell data are not as well-structured as a tetrahedron, so that the heterogeneity of quality can be expected *biologically:* a single-cell data set from a mature tissue doesn’t always have equal numbers of distinct cell types, or the cell types might have different levels of gene expression variability. What this means practically is that the distortions in a cell’s placement in the lower-dimensional representation vary in a manner that is impossible to discern “by eye”. Thus, a first step towards helping researchers use DR methods confidently is to identify *where* a dimensionally-reduced view of data is preserving high-dimensional structure and where it is not.

In Figures 4(A-C) we present lower-dimensional embeddings of the Tabula Muris marrow dataset at three different values of effective nearest neighbors in tSNE (See S3 for a discussion on how *k*_Eff_ is calculated), which is a monotonic function of its perplexity parameter. We invoke *k*_Eff_ instead of perplexity to aid with our comparisons to UMAP. The cells in these representations are colored according to the level at which the DR method was able to preserve the high-dimensional neighborhood structure relative to noise. In this color map, green is used to illustrate cells whose quality is better than 99.9% of embedded cells from a null data set. Orange then indicates cells that have a 99% chance of being better than the null, and blue indicates cells that are better represented than 90% of null cells. Pink cells are those whose neighborhoods in the lower-dimensional space are just as distorted as those generated by embedding signal-less data. As a result, this coloring allows a researcher to quantitatively understand where DR output is actually showing signal: regions of pink should not be closely interpreted since the illustrated shapes and distances are not representative of the original data. On the other hand, green regions suggest the presence of biological signal, as those parts of the embedding are unlikely to have been generated by noise. More quantitatively, a user can examine these quality levels separately, as in Figure S12, to illuminate regions that are well- or poorly-embedded.

**Figure 4:**
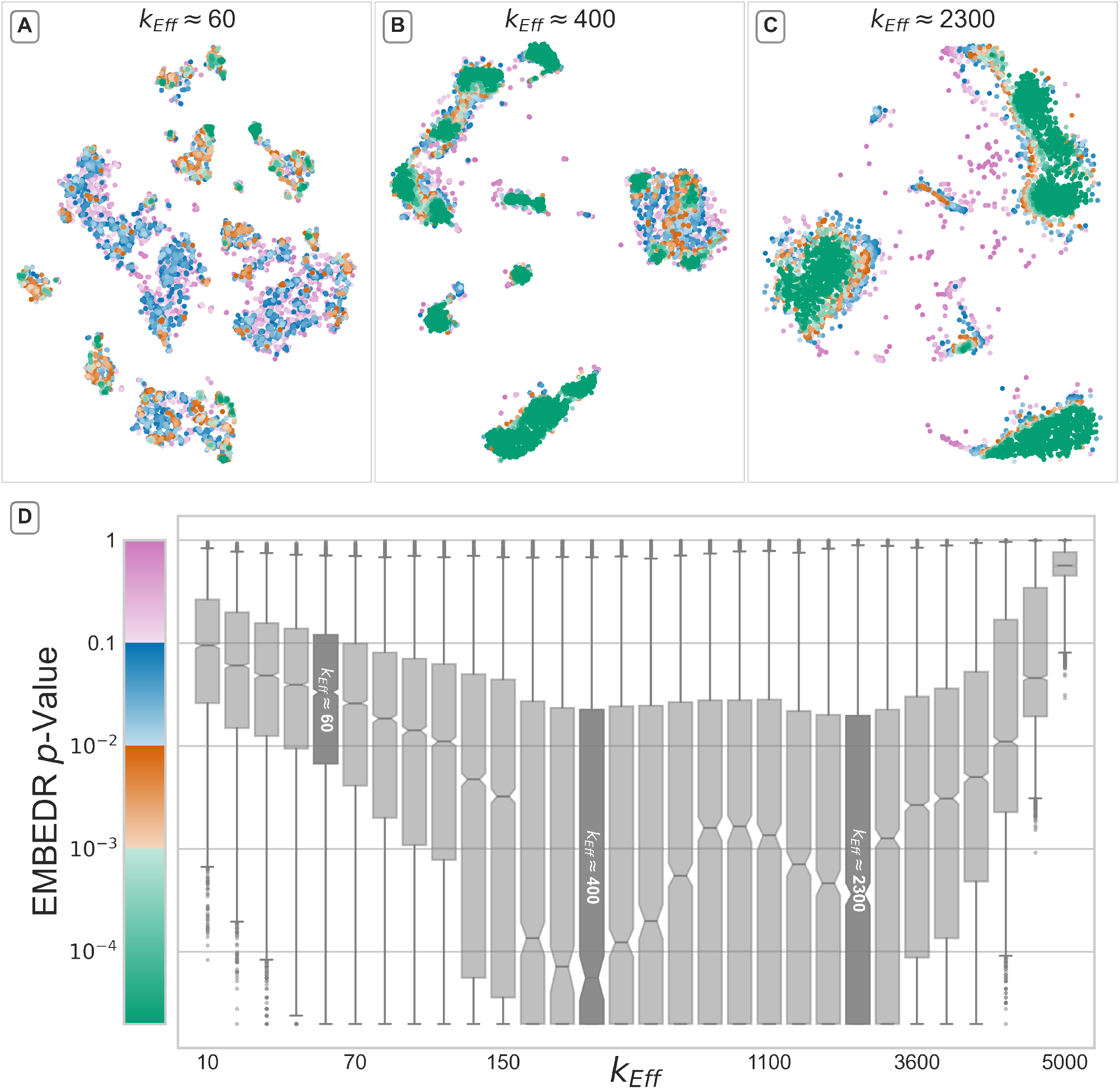
Optimizing DR Algorithm Hyperparameters Generates High-Quality Embeddings. 5037 bone marrow cells from several mice [8] were embedded with t-SNE 5 times at several values of *k*_Eff_ and the EMBEDR *p*-value was calculated using 10 null embeddings. Panels (A-C) show embeddings generated at three interesting values of *k*_Eff_; each cell is colored by the EMBEDR *p*-value. In (A), *k*_Eff_ ≈ 60 corresponds to the default t-SNE parameter in most implementations of t-SNE [22, 51]. Panel (B) shows an embedding generated using *k*_Eff_≈ 400 (`perplexity` = 250), which corresponds to the largest fraction of cells being well-represented in the lower-dimensional embedding. Similarly, panel (C) shows the results at *k*_Eff_ ≈ 2300 (`perplexity` = 1300), which corresponds to a second minimum in the *p*-values. In (D), the distributions of *p*-values are shown as box-and-whisker plots over each value of *k*_Eff_ and the median and top edge of the box plot at *k*_Eff_ ≈250 indicates that a substantial fraction of cells are best embedded at that hyperparameter value.

Generally speaking, there are some patterns worth pointing out. For example, at many values of *k*_Eff_, cells that are clustered together appear to have a similar quality of embedding — there are blue (poorly embedded) clusters and green (well embedded) clusters. We will elaborate on this further in the next section. Additionally, we observe that cells that are isolated from the center of mass of any cluster tend to be poorly embedded as well. This suggests that such cells are distinct from the rest of the data (at the scale set by *k*_Eff_) and can be classified as “outliers.” These effects can be seen in Figure S12.

It’s also worth highlighting that Figure 4(A) employs the default parameters for t-SNE, but results in a low-quality dimensionally-reduced representation of the data. 4(B) and (C) are then a potentially surprising contrast, as large portions of the data are well-represented when using hyperparameter values that are very different from common recommendations [49]. The difference in quality between these embeddings underscores the potential pitfalls of employing complex DR algorithms that require user-prescribed parameters without a quality-assessment methodology. We’ll elaborate on this more in the next section.

In this way, EMBEDR’s most immediate contribution is to provide a DR user with an intuitive map of their reduced-dimension data so that spurious structures can be separated from biological signal. A utility for generating plots like Figure 4(A-C) is included in the Python package.

### 2 EMBEDR Allows for Optimization of Algorithm Hyperparameters

As expected, Figure 4(A-C) clearly illustrates that the quality of a dimensionally-reduced view of data can vary from cell to cell across the lower-dimensional space, but Figure 4(D) shows that quality can also depend strongly on values of DR hyperparameters. In this panel, each cell’s *p*-value is summarized as box plots that change as we sweep across *k*_Eff_, the effective number of nearest neighbors used by t-SNE to place cells in two dimensions. This figure thus allows for the detection of a “globally-optimal” *k*_Eff_ based on where the largest fraction of cells are best embedded. For the Tabula Muris marrow tissue, setting *k*_Eff_ ≈ 400 corresponds to the largest fraction of minimal *p*-values, as indicated by the shaded box in Figure 4. Interestingly, *k*_Eff_ ≈ 2300 corresponds to a second dip in the *p*-values, indicating another potentially optimal hyperparameter, albeit one which is a far larger value for this parameter than is typically advised [49], even in some multiscale methods [54, 84]. This is interesting in a practical sense, as EMBEDR provides a hyperparameter tuning scheme that differs from typical heuristics.

This result also emphasizes two important considerations. First, many DR methods — t-SNE and UMAP included — have a hyperparameter that corresponds to setting the size of “neighborhoods” in the high-dimensional data. (In Supplemental S3 we show how t-SNE’s `perplexity` can be mapped to such a size, *k*_Eff_, which we use throughout this paper.) This neighborhood size then acts like a low-pass filter for electronic circuits, in that information about cells that are further than a certain “scale” is neglected. Regardless of whether a “neighborhood” is defined as a distance or as a number of nearest neighbors, however, the scale felt by the data is always mediated by the *density* of the data. What this means most directly is that the interpretation of the neighborhood size parameter must involve the size of the data set. Telling a DR method to use 15 nearest neighbors will have a very different effect when applied to a data set of 15 cells versus one with 15,000. In the former case, the effective scale is the entirety of the data, in the latter, it may be the entire data or — more likely — it may be regions that differ in size for each cell depending on the data density around that cell. As a result, these DR hyperparameters must be set and interpreted uniquely for each data set. EMBEDR’s data-sensitive statistical test means that Figure 4(D) can be constructed and interpreted consistently across data sets.

Second, the fact that large sections of cells are best embedded when t-SNE considers *k*_Eff_ ≈ 400 nearest neighbors in Figure 4(B) means that utilizing fewer neighbors for these cells may result in spurious groupings, which can be seen in the relatively poor quality of 4(A). As a result, the detection and interpretation of structures in low-dimensional representations need to account for whether the DR scale matches the “native” scale of the cells. The minima in the curve in Figure 4(D) at *k*_Eff_ ≈ 400 and 2300 mean that most cells need to consider the positions of their 400 or 2300 nearest neighbors to accurately position themselves in two dimensions, suggesting that *k*_Eff_ ≈ 400 and 2300 correspond to native scales for this data. EMBEDR facilitates this assessment by permitting comparisons between hyperparameter choices and by assessing quality locally.

The salient features of Figure 4(D) in the context of the Tabula Muris marrow dataset are preserved across the datasets we have analyzed. For example, the diaphragm tissue from the Tabula Muris Cell Atlas is analyzed in Figure S13. In all cases, EMBEDR illustrates that a) the quality of features in dimensionally-reduced data varies in a manner that is difficult to discern “by eye”, and b) the quality varies as a function of algorithmic hyperparameters and DR methods. Our ability to discern the local quality of dimensionally-reduced data results from posing the problem statistically and the generation of data-driven null hypotheses. Additionally, while it may be concerning that large portions of some DR outputs are consistent with noise, EMBEDR provides a quantitative tool with which to examine and improve these results.

### 3 EMBEDR Allows for Comparisons of Dimensionality Reduction Algorithms

Novel DR algorithms are constantly being developed or adapted, so that their incorporation into single-cell analysis requires quantitative analyses of their performance. While assessments of these methods on select case studies have been performed in [10, 11, 13, 41, 48, 49, 85], there are no results that guarantee high-performance of any of these methods on a given data set. Instead, our results and observations suggest that different methods will generate lower-dimensional embeddings with different quality for different data sets. As a result, EMBEDR’s data-driven quality assessment provides a natural tool for the comparison of DR methods applied to a common data set.

Figure 5 illustrates this approach in action, as the quality of t-SNE and UMAP embeddings of the Tabula Muris marrow data are compared side-by-side. In the top row, we show embeddings generated at t-SNE and UMAP’s default parameters, while the bottom row sets *k/k*_Eff_ based on the optima identified in Figure 4. Below each embedding, the number of cells that meet a quality threshold are indicated, showing that at default hyperparameters, neither t-SNE or UMAP generate well-matched neighborhoods for most cells. However, now the effect of optimizing t-SNE can be seen in 5(C), as nearly 50% of the cells have a neighborhood that is far more ordered than noise! When UMAP uses the same number of neighbors in 5(D), the results are improved over the defaults (5(B)), but to a slightly lesser extent than t-SNE. In Figure 5(B), the null was generated by reducing the dimensionality of the resampled data using UMAP at *k* = 15. That is, the *p*-values for each cell are determined based on how often UMAP randomly preserves structure in resampled data. Additionally, the representations in Figure 5 are colored with *p*-values generated by running t-SNE/UMAP on the data five times and on marginally resampled data ten times, so that the *p*-value indicates a *consistency* of quality as well, even though t-SNE and UMAP are stochastic and non-linear methods.

**Figure 5:**
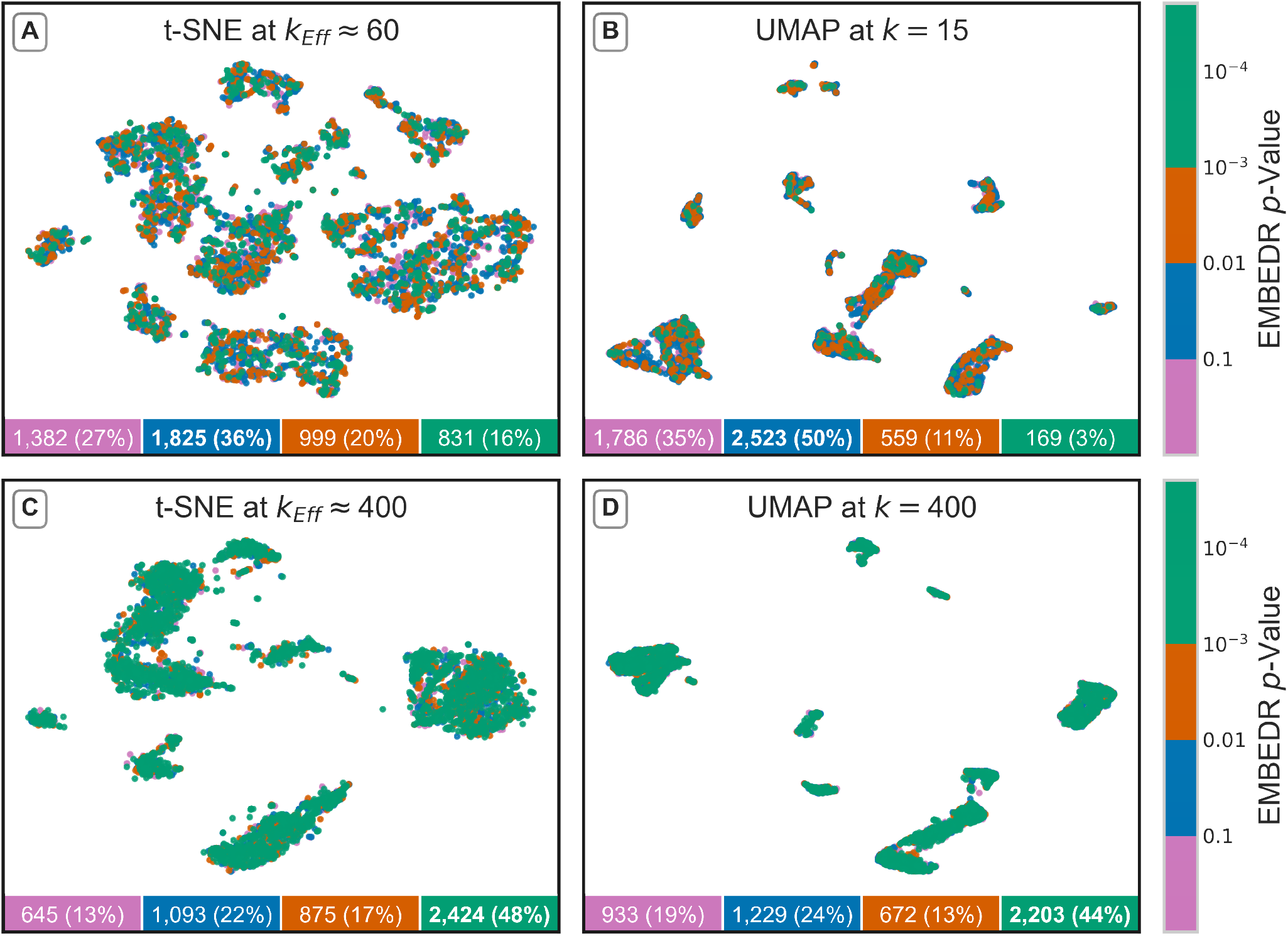
EMBEDR Facilitates Direct Comparisons of DR Methods. 5037 cells from the Tabula Muris marrow tissue [8] are embedded by t-SNE and UMAP at default (A, B) and EMBEDR-optimized (C-D) numbers of nearest neighbors. Each cell in each embedding is colored by the EMBEDR *p*-value according to the colorbars on the right. The *p*-values are calculated as in Figure 2 and Section S4 using *N*_embed_ = 5 applications of t-SNE/UMAP to the data and 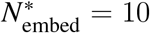 embeddings of null data. In the boxes below each panel, the number (percentage) of cells at each *p*-value threshold are shown (indicated by the corresponding color), with the threshold containing a plurality of cells shown in bold.

We emphasize that this should not be taken to mean that UMAP is not appropriate for the analysis of single-cell data, but only that t-SNE preserves structure better than UMAP in this case. We apply EMBEDR to other DR methods in Figure S14 and find similar differences in methods. Crucially, this direct, quantitative comparison of DR algorithms is an immediate consequence of our casting the quality assessment problem as a statistical problem and by generating the null hypothesis empirically.

### EMBEDR Allows for a Single-Cell Analysis of Single-Cell Data

While our results in Figures 4 and 5 show that EMBEDR can be used to push forward global analyses of DR method quality, our earlier observations that quality is heterogeneous suggest that we should be more careful and consider how embedding quality changes more *locally*. More directly, the existence of global optima in embedding quality at *k*_Eff_ ≈ 400 and 2300 does not imply that all cells are individually best embedded at those scales. Indeed, our expectation that single-cell data will contain myriad densities, cell types, and expression patterns means that we should expect to observe multiple scales in data generically. As a result, we are likely under-leveraging the information in our single-cell data by ignoring single-cell patterns.

EMBEDR provides a natural route to performing a single-cell resolution analysis of single-cell omics data as it already determines DR quality on a cell-wise basis. In Figure 6 and S15, we illustrate previously annotated cell types in the Tabula Muris marrow dataset in order to empirically demonstrate the existence of multiple scales in the data. Inspired by our observations, we propose to use a single-cell resolution analysis of single-cell data to produce a locally optimal dimensionally-reduced view of data.

**Figure 6:**
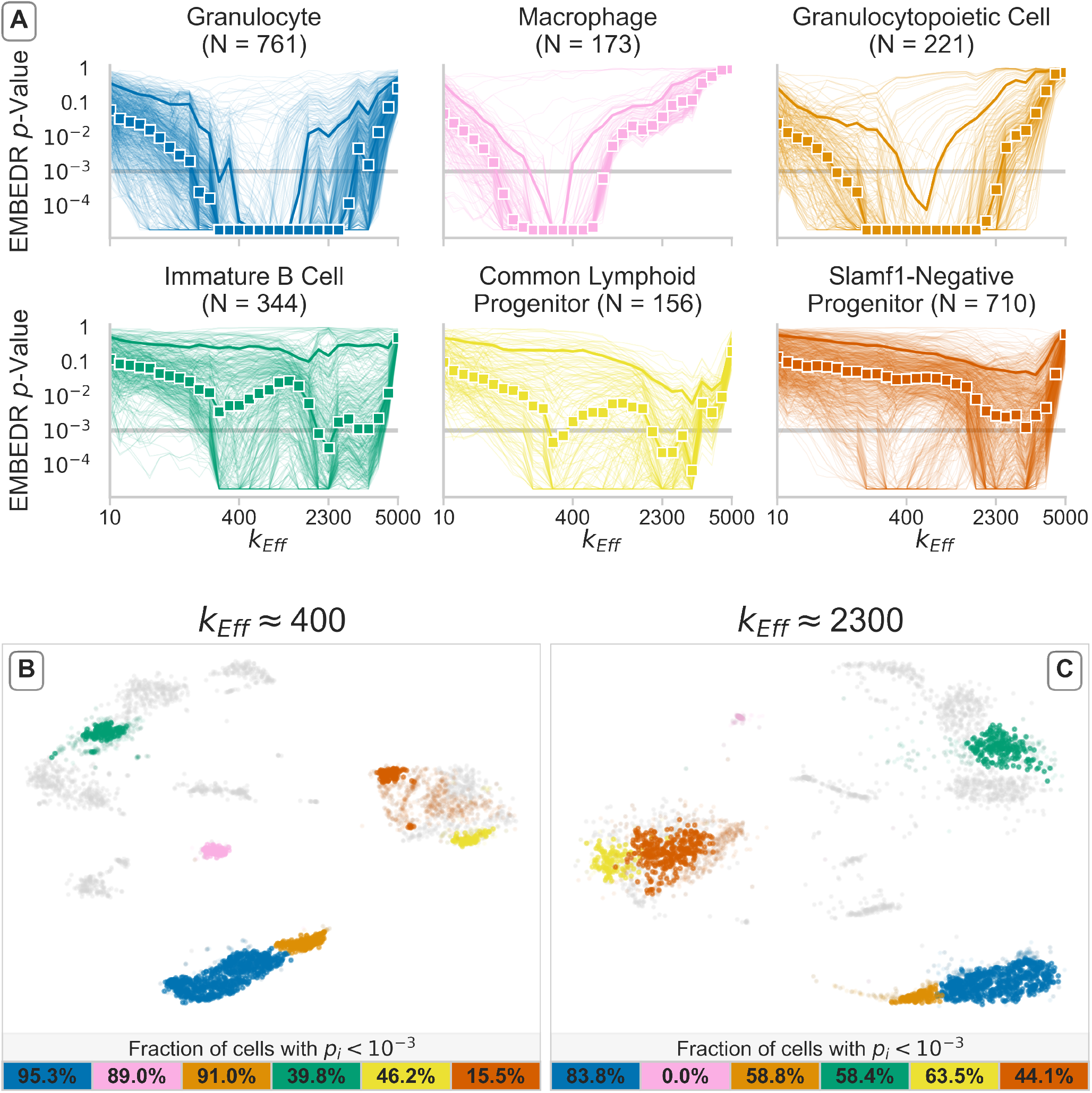
Different Cell Types are Best Embedded at a Variety of Scales. Using annotations from the Tabula Muris project [8], the embedding quality of different cell types can be examined individually across values of *k*_Eff_. In Panel (A), six identified cell types from the bone-marrow tissue are shown, where each cell with a given annotation is shown as an individual line. The colored boxes indicate the median *p*-value across all cells with that annotation, and the solid lines indicate the 90^th^ percentiles. Similar plots for all cell types are shown in Figure S15. Embeddings at *k*_Eff_ ≈ 400 and 2300 are shown in Panels (B) and (C), respectively. In (B) and (C), the cells corresponding to each cell type are highlighted with the same color as in (A). Cells with an EMBEDR *p*-value below 10^-3^ (the grey line in (A)) are opaque, while other cells with a highlighted annotation are lightly shaded. The fraction of such cells in an annotation are shown in the colored boxes below the embeddings. Other cell types are shown in grey for context.

In Figure 6(A), the EMBEDR *p*-values for cells in six select cell-types from the Tabula Muris Marrow dataset are shown as a function of *k*_Eff_. Notice that each cell’s trajectory can be followed as *k*_Eff_ increases, giving a cell-specific “spectrum”. Considering the statistics of these spectra for each cell type shows that indeed, some cell types are better represented at different scales than others. For example, macrophages (pink) appear to be well-embedded for *k*_Eff_ from 100 to 500, but the progenitor and B cells in the bottom row are best embedded in a region around *k*_Eff_ ≈ 2300. In 6(B) and (C), two examples of embeddings are different *k*_Eff_ are shown in order to illustrate the features of these spectra. In 6(B), the neighborhoods of >80% of macrophages are better structured than noise, but in 6(C) none of their neighborhoods are. The opposite happens for the progenitor and B cells: using too few neighbors results in spurious clustering and over-fracturing of these cell populations; increasing to 2300 neighbors captures that they are parts of large, diffuse regions of data space.

More generally, in the context of data sets that may contain distinct cell types, we expect this to be reflected in these spectra, as members of the same cell type may have neighborhoods at a common scale. We observe this empirically in Figure S17, where cell types with more observed cells are best represented when t-SNE uses more neighbors. If a cell is truly part of a cluster of *N* other cells, then incorporating spatial information from those *N* cells should be necessary to place that cell in an embedding. Conversely, cells from less-populous cell types are poorly placed at high *k*_Eff_ because they are being positioned using cells who are not truly their neighbors.

In this way, Figure 6 demonstrates the existence of multiple scales in the data. The differences in the spectra of cells in different cell types illustrates the sizes of different neighborhoods in the data. In this figure, the cell annotations were a given, but the relationship between EMBEDR spectra and cluster sizes (Figure S17) suggests that EMBEDR may be useful for unsupervised cluster identification. The development of such a method is beyond the scope of this work and will be pursued in the future. Instead, in Figure 7 we show how adapting t-SNE to allow for scales to be set per cell results in an improved, scale-sensitive embedding that is easily interpreted biologically.

**Figure 7:**
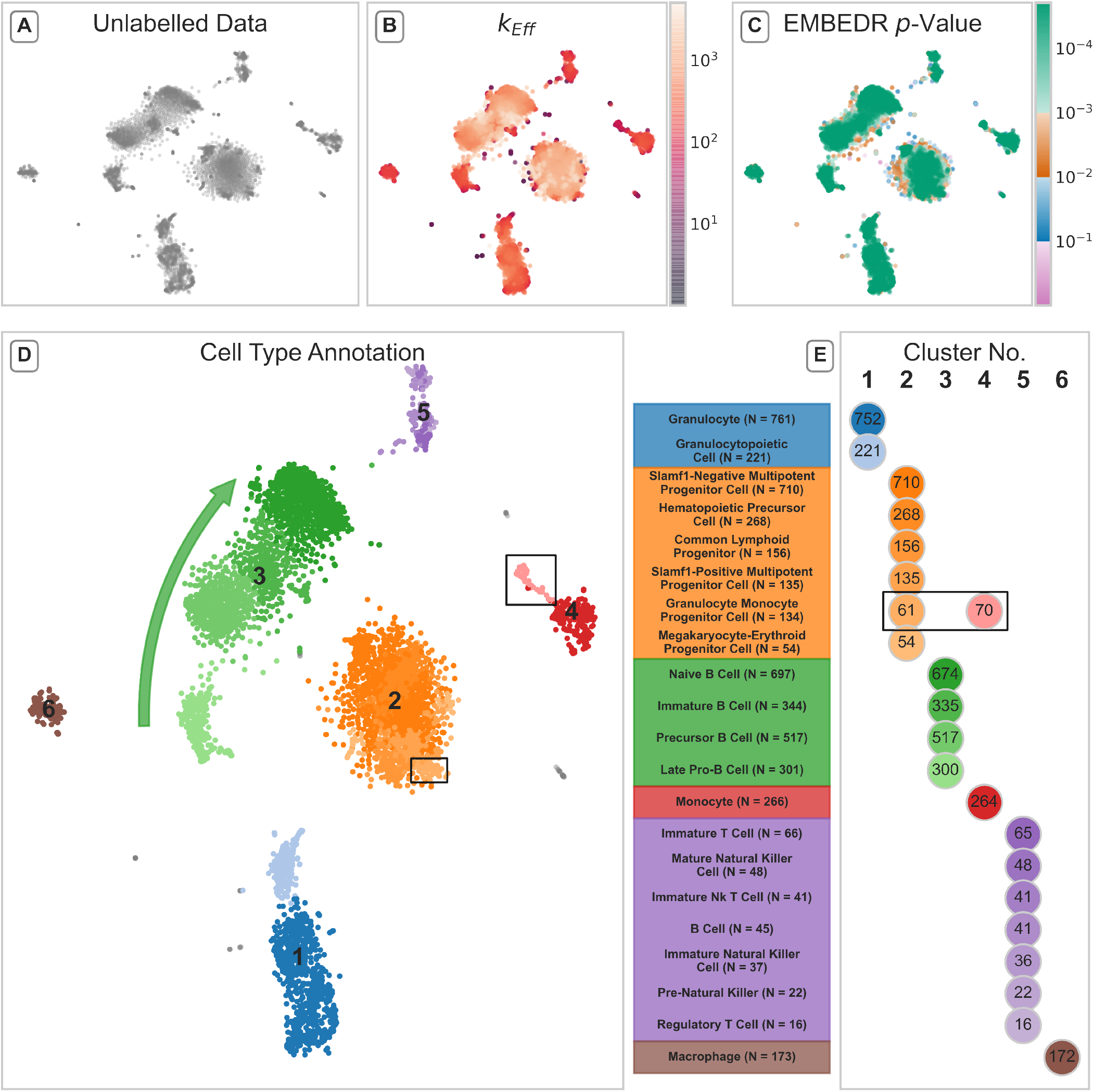
A Cell-Wise Optimized Embedding Reveals Clear Biological Signals: Adapting t-SNE to use a different scale for each sample in the Tabula Muris marrow data [8] generates a well-structured representation of the data. In (A), the unlabelled embedding is presented. To generate this embedding, the scale at which a cell’s *p*-value was minimized was used to set *k*_Eff_ for that cell. This *k*_Eff_ is shown in (B) and the minimal *p*-value is shown in (C). Applying DBSCAN with `eps` set based on the pair-wise distance distribution of cells in the embedding (specifically, the 5th percentile) detected the six indicated clusters in (D). Any Tabula Muris cell type annotation for which more than 10 cells overlapped with a DBSCAN label was given a different shade of the cluster color. These cell annotations and colors are shown in (E) as a confusion table.

Specifically, using the spectra from Figures 4 and 6 for each cell, the value for *k*_Eff_ at which each cell was best embedded was determined. These values for *k*_Eff_ were used to generate a new similarity matrix where each cell used its own “preferred” neighborhood size to determine similarities between itself and its neighbors. This similarity matrix was then used to find a representation of the data via t-SNE. The resulting embedding is shown in Figure 7. We emphasize that this representation was determined in a completely unsupervised manner.

Examination of this cell-wise optimized embedding using our established quantities in 7(B) and (C) illustrates interesting patterns. In 7(B) we see that the larger clusters were best embedded when the effective neighborhood size is large, while the smaller clusters only use *k*_Eff_≈ 100. Thus, cells that used these different scales actually ended up being clustered together, suggesting that setting these scales allows for the neighborhoods to be well-preserved and corresponds to actual signal in the data. This is reinforced by 7(C), where the minimal *p*-value of each cell is indicated, illustrating that all the large and medium-sized clusters were extremely well-embedded using cell-wise scales.

Adding labeling and annotations in 7(D) and (E) illustrates that this embedding is biologically interpretable as well. Each of the six clusters in 7(D) clearly correspond to a class of bone marrow cell types, with almost no overlap between cell annotations except for granulocyte-monocyte progenitor cells, which are indicated by black boxes. Similarly, the structure and arrangement of the clusters is biologically consistent: the annotated B cells (cluster 3, green) are all aligned according to their developmental trajectory from pro-B cells to naïve B cells. At the same time, there is no differentiation pathway in the progenitor cells (cluster 2, orange), reflecting their common multipotent state.

Looking away from the larger clusters, cells that are near the edges of clusters or between clusters tended to have a lower *k*_Eff_ and have been poorly embedded (even at their optima!). In this way, the cell-wise optimal embedding detects outlier cells in a principled manner. For example, the spread of cells between the progenitor cells in cluster 2 and the B cells in cluster 3 seem to be more reflective of low-signal cells being placed between the most heterogeneous clusters than evidence of a biological connection between the clusters. This is in contrast to other potentially spurious geometries such as the “tails” of clusters 1, 3, and 4. These regions have low *p*-values and their *k*_Eff_ suggests that their optimal scale actually involves a substantial number of nearby cells.

## Discussion

Single-cell omics offers a path towards untold biological discovery, but its high-dimensional nature and inherent stochasticity requires the careful application of dimensionality reduction algorithms in order to make progress. The promise of DR approaches to single-cell omics data is not just to gain a visual intuition for the structure of the data, but to mitigate the curse of dimensionality and perform additional downstream quantitative analyses. As of now, the state of the art in dimensionality reduction currently rests on ever-changing heuristics to a degree that limits data analysis and data-driven discovery. A researcher cannot perform a comprehensive algorithm review for each new data set, ensuring that the lack of a general approach to evaluating the quality of a DR method is preventing the community from making the most of the single-cell omics revolution. In the context of scRNA-seq, which has been the omic technology of focus in our study, cell type classification [8], lineage reconstruction [48], RNA-velocity analysis [86], and countless other approaches rely on the fidelity of dimensionally-reduced data, or are limited by their inability to confidently employ dimensionality reduction.

The statistical approach presented in this work via the EMBEDR algorithm addresses these concerns by providing a rigorous framework for the evaluation DR quality that can also reveal information about the data itself. The EMBEDR algorithm is relatively simple (Figure 2) and is available as a ready-to-use Python package. EMBEDR performs its quality assessment in a data-driven manner, meaning that it can be used to rigorously compare DR methods’ performance (Figure 5). Perhaps more importantly, EMBEDR’s local and statistical approach promises to reveal previously hidden structures in single-cell data sets while also facilitating hyperparameter optimization (Figure 4).

The EMBEDR method as proposed thus addresses the important question: “how much can I trust this dimensionally-reduced view of the data?” Embedding quality is made available as a cell-wise, interpretable *p*-value that has meaning across algorithms and data sets. This quality metric can be used to set algorithmic hyperparameters globally or locally, and can be leveraged to make inferences about the data itself. The method is robust and non-parametric.

This paper presents a broad view of the algorithm and its applications, but there are a few limitations that will require further consideration. Most practically, the code as written rests on the speed of current implementations of DR algorithms that can be chained together to generate many (null) embeddings of the same data. This is time consuming, requiring several hours to run a full scale-parameter sweep, but recently the extension of these methods to GPUs [87, 88] or quadratic rate optimization schemes [57, 89] promises drastic improvements.

The efficiency concerns also imply that there is a finite resolution to the calculated *p*-values since the null distributions are calculated empirically. This means that the number of nulls that can be embedded determines the lower-bound on the *p*-values. Other than improved computational efficiency, remedies may include theoretical work to describe the tails of these null distributions or a principled method for parameterizing the null distribution.

Moving forward, it is clear that the nature of information that EMBEDR provides can be leveraged in a variety of ways not presented in this work. Several such directions are suggested in Figure 5, where more comprehensive efforts could be undertaken to assess the quality of DR algorithms generically, such as in [40, 41]. Alternately, as suggested by Figure 6, a “spectral” view of embedding quality may provide an avenue for unsupervised clustering more directly. More simply, removing cells that never achieve a certain standard of quality may also be useful in improving traditional single-cell analyses.

Non-computationally, Figure 4 suggests that this approach may be of widespread utility in the analysis of high-dimensional biological data sets in order to detect and to assess the stability of biologically relevant structures. The ability of the method to form model-free, non-parametric scale spectra presents a new way to look at these data sets that may reveal heretofore unseen phenomena.

In all cases, high-dimensional and heterogeneous data sets such as single-cell RNA-seq require analysis techniques that account for and leverage the expected noise in the data in order to identify real biological signal. EMBEDR provides a robust statistical framework to achieve just that.

## References

1. Guo, G. et al. Resolution of Cell Fate Decisions Revealed by Single-Cell Gene Expression Analysis from Zygote to Blastocyst. Developmental Cell 18, 675–685. ISSN: 15345807 (2010).

2. Dalerba, P et al. Single-cell dissection of transcriptional heterogeneity in human colon tumors. Nature Biotechnology 29, 1120–1127. ISSN: 10870156 (2011).

3. Klein, A. M. et al. Droplet barcoding for single-cell transcriptomics applied to embryonic stem cells. Cell 161, 1187–1201. ISSN: 10974172 (May 2015).

4. Macosko, E. Z. et al. Highly parallel genome-wide expression profiling of individual cells using nanoliter droplets. Cell 161, 1202–1214. ISSN: 10974172 (2015).

5. Farrell, J. A. et al. Single-cell reconstruction of developmental trajectories during zebrafish embryogenesis. Science 360, eaar3131. ISSN: 0036-8075 (June 2018).

6. Mayer, C. et al. Developmental diversification of cortical inhibitory interneurons. Nature 555, 457–462. ISSN: 14764687 (2018).

7. Briggs, J. A. et al. The dynamics of gene expression in vertebrate embryogenesis at single-cell resolution. Science 360, eaar5780. ISSN: 0036-8075 (June 2018).

8. Schaum, N. et al. Single-cell transcriptomics of 20 mouse organs creates a Tabula Muris. Nature 562, 367–372. ISSN: 14764687 (Oct. 2018).

9. Kester, L. & van Oudenaarden, A. Single-Cell Transcriptomics Meets Lineage Tracing. Cell Stem Cell 23, 166–179. ISSN: 19345909 (Aug. 2018).

10. Hwang, B., Lee, J. H. & Bang, D. Single-cell RNA sequencing technologies and bioinformatics pipelines. Experimental and Molecular Medicine 50. ISSN: 20926413 (2018).

11. Wagner, D. E. et al. Single-cell mapping of gene expression landscapes and lineage in the zebrafish embryo. Science 360, 981–987. ISSN: 0036-8075 (June 2018).

12. Dasgupta, S., Bader, G. D. & Goyal, S. Single-Cell RNA Sequencing: A New Window into Cell Scale Dynamics. Biophysical Journal 115, 429–435. ISSN: 00063495 (Aug. 2018).

13. Grün, D. Revealing routes of cellular differentiation by single-cell RNA-seq. Current Opinion in Systems Biology 11, 9–17. ISSN: 24523100 (Oct. 2018).

14. Altman, N. & Krzywinski, M. The curse(s) of dimensionality. Nature Methods 15, 399–400. ISSN: 1548-7091 (June 2018).

15. Vallejos, C. A., Risso, D., Scialdone, A., Dudoit, S. & Marioni, J. C. Normalizing single-cell RNA sequencing data: Challenges and opportunities 2017.

16. Butler, A., Hoffman, P., Smibert, P., Papalexi, E. & Satija, R. Integrating single-cell transcriptomic data across different conditions, technologies, and species. Nature Biotechnology 36, 411–420. ISSN: 15461696 (June 2018).

17. Gong, W., Kwak, I. Y., Pota, P., Koyano-Nakagawa, N. & Garry, D. J. DrImpute: Imputing dropout events in single cell RNA sequencing data. BMC Bioinformatics 19. ISSN: 14712105 (2018).

18. Hafemeister, C. & Satija, R. Normalization and variance stabilization of single-cell RNA-seq data using regularized negative binomial regression. Genome Biology 20, 296. ISSN: 1474-760X (Dec. 2019).

19. Huang, M. et al. SAVER: gene expression recovery for single-cell RNA sequencing. Nature Methods 15, 539–542. ISSN: 1548-7091 (July 2018).

20. Lähnemann, D. et al. Eleven grand challenges in single-cell data science. Genome Biology 21, 31. ISSN: 1474-760X (Dec. 2020).

21. Jollife, I. T. & Cadima, J. Principal component analysis: A review and recent developments. Philosophical Transactions of the Royal Society A: Mathematical, Physical and Engineering Sciences 374. ISSN: 1364503X (2016).

22. Van Der Maaten, L. & Hinton, G. Visualizing Data using t-SNE. Journal of Machine Learning Research 9, 2579–2605. ISSN: 15729338 (2008).

23. McInnes, L., Healy, J. & Melville, J. UMAP: Uniform Manifold Approximation and Projection for Dimension Reduction. arXiv (Feb. 2018).

24. Kohonen, T. Self-organized formation of topologically correct feature maps. Biological Cybernetics 43, 59–69. ISSN: 0340-1200 (1982).

25. Schölkopf, B., Smola, A. & Müller, K.-R. Nonlinear Component Analysis as a Kernel Eigenvalue Problem. Neural Computation 10, 1299–1319. ISSN: 0899-7667 (July 1998).

26. Tenenbaum, J. B., de Silva, V. & Langford, J. C. A Global Geometric Framework for Nonlinear Dimensionality Reduction. Science 290, 2319–2323. ISSN: 00368075 (Dec. 2000).

27. Roweis, S. T. Nonlinear Dimensionality Reduction by Locally Linear Embedding. Science 290, 2323–2326. ISSN: 00368075 (Dec. 2000).

28. Belkin, M. & Niyogi, P. Laplacian Eigenmaps for Dimensionality Reduction and Data Representation. Neural Computation 15, 1373–1396. ISSN: 0899-7667 (June 2003).

29. Chen, M. et al. The Bayesian Elastic Net: Classifying Multi-Task Gene-Expression Data (2009).

30. Venna, J., Kaski, S., Aidos, H., Nybo, K. & Peltonen, J. Information retrieval perspective to nonlinear dimensionality reduction for data visualization. Journal of Machine Learning Research 11, 451–490. ISSN: 15324435 (2010).

31. Joia, P., Paulovich, F. V., Coimbra, D., Cuminato, J. A. & Nonato, L. G. Local Affine Multidimensional Projection. IEEE Transactions on Visualization and Computer Graphics 17, 2563–2571. ISSN: 1077-2626 (Dec. 2011).

32. Najim, S. A. & Lim, I. S. Trustworthy dimension reduction for visualization different data sets. Information Sciences 278, 206–220. ISSN: 00200255 (Sept. 2014).

33. Wang, B., Zhu, J., Pierson, E., Ramazzotti, D. & Batzoglou, S. Visualization and analysis of single-cell rna-seq data by kernel-based similarity learning. Nature Methods 14, 414–416. ISSN: 15487105 (2017).

34. Risso, D., Perraudeau, F., Gribkova, S., Dudoit, S. & Vert, J. P. A general and flexible method for signal extraction from single-cell RNA-seq data. Nature Communications 9. ISSN: 20411723 (2018).

35. Wu, Y., Tamayo, P. & Zhang, K. Visualizing and Interpreting Single-Cell Gene Expression Datasets with Similarity Weighted Nonnegative Embedding. Cell Systems 7, 656–666. ISSN: 24054712 (Dec. 2018).

36. Tarashansky, A. J., Xue, Y., Li, P., Quake, S. R. & Wang, B. Self-assembling manifolds in single-cell RNA sequencing data. eLife 8, 1–29. ISSN: 2050084X (Sept. 2019).

37. Moon, K. R. et al. Visualizing structure and transitions in high-dimensional biological data. Nature Biotechnology 37, 1482–1492. ISSN: 15461696 (2019).

38. Eraslan, G., Simon, L. M., Mircea, M., Mueller, N. S. & Theis, F. J. Single-cell RNA-seq denoising using a deep count autoencoder. Nature Communications 10, 390. ISSN: 2041-1723 (Dec. 2019).

39. Van Der Maaten, L. J. P., Postma, E. O. & Van Den Herik, H. J. Dimensionality Reduction: A Comparative Review. Journal of Machine Learning Research 10, 1–41. ISSN: 0169328X (2009).

40. Gracia, A., González, S., Robles, V. & Menasalvas, E. A methodology to compare Dimensionality Reduction algorithms in terms of loss of quality. Information Sciences 270, 1–27. ISSN: 00200255 (June 2014).

41. Espadoto, M., Martins, R. M., Kerren, A., Hirata, N. S. T. & Telea, A. C. Towards a Quantitative Survey of Dimension Reduction Techniques. IEEE Transactions on Visualization and Computer Graphics X, 1–1. ISSN: 1077-2626 (2019).

42. Fanaee-T, H. & Thoresen, M. Performance evaluation of methods for integrative dimension reduction. Information Sciences 493, 105–119. ISSN: 00200255 (Aug. 2019).

43. Gracia, A., González, S., Robles, V., Menasalvas, E. & Von Landesberger, T. New insights into the suitability of the third dimension for visualizing multivariate/multidimensional data: A study based on loss of quality quantification. Information Visualization 15, 3–30. ISSN: 14738724 (Jan. 2016).

44. Lui, K. Y. C., Ding, G. W., Huang, R. & McCann, R. J. Dimensionality Reduction has Quantifiable Imperfections: Two Geometric Bounds. Advances in Neural Information Processing Systems 2018-Decem, 8453–8463. ISSN: 10495258 (Oct. 2018).

45. Aupetit, M. Visualizing distortions and recovering topology in continuous projection techniques. Neurocomputing 70, 1304–1330. ISSN: 09252312 (Mar. 2007).

46. Mokbel, B., Lueks, W., Gisbrecht, A. & Hammer, B. Visualizing the quality of dimensionality reduction. Neurocomputing 112, 109–123. ISSN: 0925-2312 (July 2013).

47. Colange, B., Vuillon, L., Lespinats, S. & Dutykh, D. Interpreting Distortions in Dimensionality Reduction by Superimposing Neighbourhood Graphs in 2019 IEEE Visualization Conference (VIS) (IEEE, Oct. 2019), 211–215. ISBN: 978-1-7281-4941-7.

48. Herring, C. A., Chen, B., McKinley, E. T. & Lau, K. S. Single-Cell Computational Strategies for Lineage Reconstruction in Tissue Systems. Cmgh 5, 539–548. ISSN: 2352345X (2018).

49. Kobak, D. & Berens, P. The art of using t-SNE for single-cell transcriptomics. Nature Communications 10, 5416. ISSN: 2041-1723 (Dec. 2019).

50. France, S. L. & Akkucuk, U. A Review, Framework and R toolkit for Exploring, Evaluating, and Comparing Visualizations (Feb. 2019).

51. Poličar, P., Stražar, M. & Zupan, B. openTSNE: a modular Python library for t-SNE dimensionality reduction and embedding. bioRxiv, 1–2 (2019).

52. Lee, J. A., Peluffo-Ordóñez, D. H. & Verleysen, M. Multiscale stochastic neighbor embedding: Towards parameter-free dimensionality reduction in 22nd European Symposium on Artificial Neural Networks, Computational Intelligence and Machine Learning, ESANN 2014 - Proceedings (2014), 177–182. ISBN: 9782874190957.

53. Cao, Y. & Wang, L. Automatic Selection of t-SNE Perplexity. arXiv (Aug. 2017).

54. Bodt, C. D., Mulders, D., Verleysen, M. & Lee, J. A. Perplexity-free t-SNE and twice Student tt -SNE in European Symposium on Artificial Neural Networks (Bruges, Belgium, 2018). ISBN: 978-287587047-6.

55. Linderman, G. C., Rachh, M., Hoskins, J. G., Steinerberger, S. & Kluger, Y. Fast interpolation-based t-SNE for improved visualization of single-cell RNA-seq data. Nature Methods 16, 243–245. ISSN: 1548-7091 (Mar. 2019).

56. Aliverti, E. et al. Projected t-SNE for batch correction. Bioinformatics 36 (ed Wren, J.) 3522–3527. ISSN: 1367-4803 (June 2020).

57. Häkkinen, A. et al. qSNE: Quadratic rate t-SNE optimizer with automatic parameter tuning for large data sets. Bioinformatics, 1–7. ISSN: 1367-4803 (2020).

58. Belkina, A. C. et al. Automated optimized parameters for T-distributed stochastic neighbor embedding improve visualization and analysis of large datasets. Nature Communications 10, 5415. ISSN: 2041-1723 (Dec. 2019).

59. Lee, J. A. & Verleysen, M. Quality assessment of dimensionality reduction: Rank-based criteria. Neurocomputing 72, 1431–1443. ISSN: 09252312 (Mar. 2009).

60. Venna, J. & Kaski, S. in Lecture Notes in Computer Science (including subseries Lecture Notes in Artificial Intelligence and Lecture Notes in Bioinformatics) September, 485–491 (2001). ISBN: 3540424865.

61. France, S. & Carroll, D. in Machine Learning and Data Mining in Pattern Recognition 499–517 (Springer Berlin Heidelberg, Berlin, Heidelberg, 2007).

62. Lee, J. A. & Verleysen, M. Quality assessment of nonlinear dimensionality reduction based on {K}-ary neighborhoods in JMLR: Workshop and conference proceedings 4 (2008), 21–35.

63. Goldberg, Y. & Ritov, Y. Local procrustes for manifold embedding: a measure of embedding quality and embedding algorithms. Machine Learning 77, 1–25. ISSN: 0885-6125 (Oct. 2009).

64. Lee, A. Circular data. Wiley Interdisciplinary Reviews: Computational Statistics 2, 477–486. ISSN: 19395108 (2010).

65. Meng, D., Leung, Y. & Xu, Z. A new quality assessment criterion for nonlinear dimensionality reduction. Neurocomputing 74, 941–948. ISSN: 09252312 (Feb. 2011).

66. Zhang, P., Ren, Y. & Zhang, B. A new embedding quality assessment method for manifold learning. Neurocomputing 97, 251–266. ISSN: 09252312 (Nov. 2012).

67. Paul, R. & Chalup, S. K. A study on validating non-linear dimensionality reduction using persistent homology. Pattern Recognition Letters 100, 160–166 (Dec. 2017).

68. Heiser, C. N. & Lau, K. S. A Quantitative Framework for Evaluating Single-Cell Data Structure Preservation by Dimensionality Reduction Techniques. Cell Reports 31, 107576. ISSN: 22111247 (2020).

69. Kaski, S. et al. Trustworthiness and metrics in visualizing similarity of gene expression. BMC Bioinformatics 4. ISSN: 14712105 (2003).

70. Lespinats, S. & Aupetit, M. CheckViz: Sanity Check and Topological Clues for Linear and Non-Linear Mappings. Computer Graphics Forum 30, 113–125. ISSN: 01677055 (Mar. 2011).

71. Schreck, T., von Landesberger, T. & Bremm, S. Techniques for precision-based visual analysis of projected data in Visualization and Data Analysis 2010 (eds Park, J., Hao, M. C., Wong, P. C. & Chen, C.) 7530 (Jan. 2010), 75300E. ISBN: 9780819479235.

72. Martins, R. M., Minghim, R. & Telea, A. C. Explaining neighborhood preservation for multidimensional projections. Computer Graphics and Visual Computing, CGVC 2015, 7–14 (2015).

73. Rieck, B. & Leitte, H. Persistent Homology for the Evaluation of Dimensionality Reduction Schemes. Computer Graphics Forum 34, 431–440. ISSN: 01677055 (June 2015).

74. Rieck, B. & Leitte, H. in Topological Methods in Data Analysis and Visualization IV (eds Carr, H., Garth, C. & Weinkauf, T.) 103–117 (Springer International Publishing, Cham, 2017). ISBN: 978-3-319-44684-4.

75. Martins, R. M., Coimbra, D. B., Minghim, R. & Telea, A. Visual analysis of dimensionality reduction quality for parameterized projections. Computers & Graphics 41, 26–42. ISSN: 00978493 (June 2014).

76. Kullback, S. & Leibler, R. A. On Information and Sufficiency. The Annals of Mathematical Statistics 22, 79–86. ISSN: 0003-4851 (Mar. 1951).

77. Lee, J. A., Renard, E., Bernard, G., Dupont, P. & Verleysen, M. Type 1 and 2 mixtures of Kullback-Leibler divergences as cost functions in dimensionality reduction based on similarity preservation. Neurocomputing 112, 92–108. ISSN: 09252312 (2013).

78. Halabi, N., Rivoire, O., Leibler, S. & Ranganathan, R. Protein Sectors: Evolutionary Units of Three-Dimensional Structure. Cell 138, 774–786. ISSN: 1097-4172 (Aug. 2009).

79. Plerou, V. et al. A Random Matrix Approach to Cross-Correlations in Financial Data. Physical Review E 65, 066126. ISSN: 1063-651X (Aug. 2001).

80. Aparicio, L., Bordyuh, M., Blumberg, A. J. & Rabadan, R. A Random Matrix Theory Approach to Denoise Single-Cell Data. Patterns 1, 100035. ISSN: 26663899 (June 2020).

81. Loughin, T. M. A systematic comparison of methods for combining *p*-values from independent tests. Computational Statistics and Data Analysis 47, 467–485. ISSN: 01679473 (2004).

82. Cousins, R. D. Annotated Bibliography of Some Papers on Combining Significances or *p*-values. arXiv (May 2007).

83. Heard, N. & Rubin-Delanchy, P. Choosing Between Methods of Combining *p*-values (July 2017).

84. Lee, J. A., Peluffo-Ordóñez, D. H. & Verleysen, M. Multi-scale similarities in stochastic neighbour embedding: Reducing dimensionality while preserving both local and global structure. Neurocomputing 169, 246–261. ISSN: 09252312 (Dec. 2015).

85. Gisbrecht, A. & Hammer, B. Data visualization by nonlinear dimensionality reduction. Wiley In-terdisciplinary Reviews: Data Mining and Knowledge Discovery 5, 51–73. ISSN: 19424787 (Mar. 2015).

86. La Manno, G. et al. RNA velocity of single cells. Nature 560, 494–498. ISSN: 0028-0836 (Aug. 2018).

87. Chan, D. M., Rao, R., Huang, F. & Canny, J. F. T-SNE-CUDA: GPU-Accelerated T-SNE and its Applications to Modern Data in 2018 30th International Symposium on Computer Architecture and High Performance Computing (SBAC-PAD) (IEEE, Sept. 2018), 330–338. ISBN: 978-1-5386-7769-8.

88. Agrawal, A., Ali, A. & Boyd, S. Minimum-Distortion Embedding tech. rep. (2021).

89. De Bodt, C., Mulders, D., Verleysen, M. & Lee, J. A. Fast Multiscale Neighbor Embedding. IEEE Transactions on Neural Networks and Learning Systems, 1–15. ISSN: 2162-237X (2020).

90. Kobak, D., Linderman, G., Steinerberger, S., Kluger, Y. & Berens, P. Heavy-Tailed Kernels Reveal a Finer Cluster Structure in t-SNE Visualisations in Machine Learning and Knowledge Discovery in Databases (eds Brefeld, U. et al.) 11906 LNAI (Springer International Publishing, Cham, Feb. 2020), 124–139. ISBN: 978-3-030-46150-8.

91. Narayan, A., Berger, B. & Cho, H. Density-Preserving Data Visualization Unveils Dynamic Patterns of Single-Cell Transcriptomic Variability.

92. Vovk, V. & Wang, R. Combining p-values via averaging. Biometrika 107, 791–808. ISSN: 0006-3444 (Dec. 2020).

93. LeCun, Y., Bottou, L., Bengio, Y. & Haffner, P Gradient-based learning applied to document recognition. Proceedings of the IEEE 86, 2278–2323. ISSN: 00189219 (1998).

94. Li, P., Hastie, T. J. & Church, K. W. Very sparse random projections in Proceedings of the 12th ACM SIGKDD international conference on Knowledge discovery and data mining - KDD ‘06 2006 (ACM Press, New York, New York, USA, 2006), 287. ISBN: 1595933395.

