## Supplemental for "EMBEDR: Distinguishing Signal from Noise in Single-Cell Omics Data"

### Supplemental Materials

#### S1 Calculating a Cell-Wise Kullback-Liebler Divergence

When defining a "neighborhood" of a sample in a space, it is natural to use either a rank-based radius, such as the number of nearest neighbors, $k$, or a distance-based radius, such as a specific Euclidean distance, $d_r$. Unfortunately, both these methods for specifying scale are sensitive to the *local density* of samples, and if the density of samples in a data set varies, then the relationship between $d_r$ and $k$ will vary as well. To circumvent this, t-SNE, and SNE methods more generally, use "affinities" between samples where the scale of the "affinity" can be set locally, for each sample, so that all neighborhoods preserve some other quantity, such as entropy [22, 52, 54, 77, 89–91].

In t-SNE [22], the "affinity" between two samples, $\vec{x}_i$ and $\vec{x}_j$, is based on the Gaussian probability that cell $j$ is a distance, $d_{ij} = \left|\vec{x}_i - \vec{x}_j\right|$, away. That is,

$$p_{j|i} = \mathcal{N}\left(d_{ij}; 0, \sigma_i\right) \tag{1}$$

The width of this Gaussian kernel, $\sigma_i$, is specified by the `perplexity` parameter. Specifically,

$$\texttt{perplexity} = e^{H_i}, \text{ where } H_i = -\sum_j p_{j|i} \log\left(p_{j|i}\right) \tag{2}$$

is the *entropy* of the Gaussian kernel. $\sigma_i$ is set by performing a binary search over possible widths until the distribution has the required entropy.

SNE methods then also calculate an affinity between samples in the low-dimensional space, indicated $q_{j|i}$. In t-SNE,

$$q_{j|i} = q_{ij} = \frac{\left(1 + \left|\vec{y}_i - \vec{y}_j\right|^2\right)^{-1}}{\sum_{k \neq l}\left(1 + |\vec{y}_k - \vec{y}_l|^2\right)^{-1}}, \tag{3}$$

i.e. the low-dimensional affinities are calculated using a Student's $t$-distribution with degrees of freedom, $\nu = 1$.

t-SNE then uses gradient descent to minimize the Kullback-Liebler divergence, $\mathcal{D}_{KL}$, between the high-dimensional affinity matrix, $P = \left\{p_{ij}\right\}$, and the low-dimensional affinity matrix, $Q = \left\{q_{ij}\right\}$. (t-SNE and other methods often *symmetrize* the affinity matrices so that $p_{ij} \propto p_{j|i} + p_{i|j}$.) In calculating $\mathcal{D}_{KL}(P||Q)$, it is assumed that $P$ and $Q$ are probability distributions, so it is necessary to normalize so that $\sum_{i,j} p_{ij} = \sum_{i,j} q_{ij} = 1$.

In this work, we note that we can maintain sample-wise affinities by *row-normalizing* $P$ and $Q$ so that

$$\hat{p}_{j|i} = \frac{p_{j|i}}{\sum_{k \neq i} p_{k|i}} \qquad \text{and} \qquad \hat{q}_{ij} = \frac{\left(1 + \left|\vec{y}_i - \vec{y}_j\right|^2\right)^{-1}}{\sum_{k \neq i}\left(1 + |\vec{y}_i - \vec{y}_k|^2\right)^{-1}}. \tag{4}$$

We then define $P_i = \left\{\hat{p}_{j|i}\right\}_{j=1,\ldots,N}$ and $Q_i = \left\{\hat{q}_{ij}\right\}_{j=1,\ldots,N}$ so that we can calculate the sample-wise divergence between kernels in the high- and low-dimensional space:

$$\mathcal{D}_{KL}^i = \mathcal{D}_{KL}(P_i||Q_i) = \sum_{j=1}^{N} \hat{p}_{j|i} \log\left(\frac{\hat{p}_{j|i}}{\hat{q}_{ij}}\right). \tag{5}$$

661 Throughout the paper, we set $\mathcal{D}_{KL}^i = EES_i$ to be the cell-wise quality metric.

#### 662 S2 t-SNE's Global $D_{KL}$ can be broken into a sum of cell-wise $D_{KL}$

Here we show how the t-SNE Kulback-Leibler divergence can be separated into cell-wise (point-wise) measures. We start with the t-SNE K.L. definition,

$$\mathcal{D}(P||Q) = \sum_{ij} p_{ij} \ln\left(\frac{p_{ij}}{q_{ij}}\right).$$

We next define the row-by-row marginals and conditional probabilities:

$$\hat{p}_{j|i} = \frac{p_{ij}}{p_i} \quad \text{with} \quad p_i = \sum_j p_{ij} \quad \text{and} \quad \hat{q}_{j|i} = \frac{q_{ij}}{q_i} \quad \text{with} \quad q_i = \sum_j q_{ij}\,.$$

The $\hat{p}_{j|i}$ and $\hat{q}_{j|i}$ here are conditional probabilities, *not* the unsymmetrized probabilities defined in the t-SNE algorithm [22] (see Section S1). Note also that

$$p_{ij} = \hat{p}_{j|i} p_i \quad \text{and} \quad q_{ij} = \hat{q}_{j|i} q_i\,.$$

Substituting the latter into the K.L. divergence gives

$$\begin{aligned}\mathcal{D}(P||Q) &= \sum_{ij} \hat{p}_{j|i} p_i \ln\left(\frac{\hat{p}_{j|i} p_i}{\hat{q}_{j|i} q_i}\right) = \sum_{ij} \hat{p}_{j|i} p_i \left[\ln\left(\frac{p_i}{q_i}\right) + \ln\left(\frac{\hat{p}_{j|i}}{\hat{q}_{j|i}}\right)\right] \\ &= \sum_i p_i \ln\left(\frac{p_i}{q_i}\right) + \sum_i p_i \sum_j \hat{p}_{j|i} \ln\left(\frac{\hat{p}_{j|i}}{\hat{q}_{j|i}}\right)\end{aligned}$$

where we have used $\sum_j \hat{p}_{j|i} = 1$. The above can also be written

$$\mathcal{D}(P||Q) = \sum_i p_i \left[\ln\left(\frac{p_i}{q_i}\right) + \sum_j \hat{p}_{j|i} \ln\left(\frac{\hat{p}_{j|i}}{\hat{q}_{j|i}}\right)\right].$$

If we know we are considering only a particular row, then we normalize so that the $p_i$ and $q_i$ for that row become 1, and one can define the row-specific K.L. divergence as

$$\mathcal{D}_{KL}^i(P_i||Q_i) = \sum_j \hat{p}_{j|i} \ln\left(\frac{\hat{p}_{j|i}}{\hat{q}_{j|i}}\right).$$

663 The total K.L. divergence, if one requires it, is then the sum $\sum_i p_i \left[\ln(p_i/q_i) + \mathcal{D}_{KL}^i\right]$.

It's worth noting that UMAP minimizes a similar quantity in its optimization, the *cross-entropy*, which is the sum of the Kullback-Liebler Divergence and the entropy of $P$, $H(P)$. As a result, $\mathcal{D}^{i}_{KL}$ is a natural quantity for both these methods.

### S3 Interpreting t-SNE's `perplexity` parameter as an "effective" number of nearest neighbors

Despite `perplexity` being an innovative and useful parameterization of the problem — the t-SNE methodology ensures that all neighborhoods contain similar amounts of *information*[76] — it is notoriously difficult to set and interpret [49, 52, 54]. We note that heuristically, `perplexity` corresponds to setting a *scale* in the data in that generally the size of the Gaussian kernel will need to grow with `perplexity` in order to add the requisite entropy. In this section, we formalize a process by which `perplexity` can be unambiguously converted into an "effective number of nearest neighbors", $k_{\text{Eff}}$, for a given data set.

Before we define our heuristic, we note that contrary to popular belief, `perplexity` is a parameter that can take on any non-negative value, not just integers. However, there are some useful bounds that we can find by considering Equation 2. First, if we consider an affinity distribution on $N$ samples, its entropy is maximized if the distribution is uniform ($p_{j|i} = 1/N$ for all $j = 1, \ldots, N$). In such a case, the corresponding perplexity can be found to be

$$\texttt{perplexity}_{\text{max}} = e^{-\sum \frac{1}{N}\log\left(\frac{1}{N}\right)} = e^{-\log\left(\frac{1}{N}\right)} = N. \tag{6}$$

Setting `perplexity`$= N$ thus corresponds to giving all samples an equal weight so that t-SNE becomes similar to a spectral method [55].

In the other extreme, if all the affinity is localized to one neighbor, $k$, so that $p_{k|i} = 1$ and $p_{j|i} = 0$ for all $j \neq k$, then we get

$$\texttt{perplexity}_{\text{min}} = e^{-(1)\log(1) - \sum_{j\neq k}(0)\log(0)} = e^{0} = 1. \tag{7}$$

That is, such an affinity distribution has no entropy ($H_i = 0$). Thus, the `perplexity` parameter is constrained to take on values between 1 and $N$. This is notable because it means that the interpretation of a value of `perplexity` depends on the size of the data set! This observation motivates our exploration of embedding quality for `perplexity` values that are much larger than are typically examined (`perplexity`$\rightarrow N$).

Once a value for `perplexity` has been specified, the size of the Gaussian kernel, $\sigma_i$, is fixed via a binary search as in [22]. This kernel is fixed over $k$ neighbors (typically $k = 3\times$`perplexity`), but that does not mean that all $k$ neighbors contribute meaningfully to the affinity distribution. As an example, consider Figure S6, where the affinities (kernel probabilities) for 90 nearest neighbors of a sample point in the Tabula Muris Marrow data set [8] are shown as a function of their distance to the sample point and their rank distance to the sample point. Note that while all neighbors have nonzero affinity with the sample point, the nearer neighbors have orders of magnitude more weight than the more distant ones. This suggests that the width of these kernels can be converted into a number of nearest neighbors that contribute "meaningully" to the distribution, i.e. an "effective number of nearest neighbors," $k_{\text{Eff}}$.

In Figure S6, we illustrate three potential markers that can be converted into a $k_{\text{Eff}}$:

1. a threshold based on the kernel width $\sigma_i$ (green),
2. a global threshold on the affinities (red),

3. or a local threshold based on the ratio of affinities to the affinity of the nearest neighbor (purple).

Because the kernel is uniquely fixed based on each cell's neighbors, setting a global threshold (red) or a kernel-based threshold (green) won't measure the number of cells that are meaningfully contributing to an affinity distribution. That is, the exact spacing of the nearest neighbors changes the interpretation of those thresholds so that they are hard to generalize across all cells. On the other hand, the local threshold (purple) responds both to the neighbor spacing and the distribution's size in a consistent manner. As a result, we set a cell's effective nearest neighbors to be those cells that have an affinity above some percentage of that of the nearest neighbors.

$$k_{\mathrm{Eff}}{}^{i} = \left| j; p_{j|i} \geq \alpha \max_{j \neq i} p_{j|i} \right| \tag{8}$$

We then need to choose $\alpha$ in such a way that corresponds to our semantic understanding of "effective neighbors". Using Figures S7 and S8, we find that $\alpha = 0.01$ provides a balance between capturing most of the neighborhood information (the relative change in embedding quality is $< 10\%$) while also not using all available neighbors. (Setting $\alpha$ too small simply says that all neighbors are effective so that there is no change in quality.)

Once we have found each cell's effective neighborhood size for each value of `perplexity`, we can use Figure S9 to establish an *average* relationship between $k_{\mathrm{Eff}}$ and `perplexity` by interpolating between the median $k_{\mathrm{Eff}}{}^{i}$ at chosen values of `perplexity`. In this way, we can use $k_{\mathrm{Eff}}$ to indicate the scale of the neighborhoods as opposed to the more inpenetrable `perplexity` parameter.

It's worth noting that this process is only a heuristic to facilitate the interpretation of analyses that vary `perplexity`. Using this heuristic allows us to compare DR methods such as t-SNE and UMAP in a more meaningful manner, as we do in Figure 5. This heuristic is also *data-dependent*; the relationship between `perplexity` and $k_{\mathrm{Eff}}$ depends on the dimensionality, density, and structure of the data. However, we expect the relationship to be monotonic for most data, making it a useful heuristic for interpreting t-SNE analyses.

### S4 Calculating a Consensus $p$-Value

The last step of the EMBEDR algorithm involves comparing each cell's embedding quality, $EES_i$ to the null distribution for $EES_i^*$ to calculate an empirical $p$-value for the likelihood that the null data could generate such a quality score. We noted that many DR algorithms are stochastic or non-linear, so that a cell's embedding quality may vary between runs of these algorithms (see Figure S10). As a result, each run of a stochastic DR algorithm will generate a new EMBEDR $p$-value for each cell. However, for visualization and analysis purposes, it is more useful to have a single quantity that captures the quality of a cell's position across several embeddings.

To summarize these multiple $p$-values for each cell, we consider each embedding to be a unique, dependent measurement of the quality statistic. There are then several possible methods with which to consolidate these measurements [82, 83, 92], and we find that the simplest is to simply average the $p$-values across embeddings.

Besides being computationally simple, an average of $N_{\mathrm{embed}}$ $p$-values can also be understood to be an estimate of the likelihood that a random variable $X$ is greater than another random variable $Y$. To see this, we can write

$$P(X > Y) = \iint_{x>y} p_X(x) p_Y(y) dx dy = \int_{-\infty}^{\infty} \int_{-\infty}^{x} p_Y(y) p_X(x) dy dx = \int_{-\infty}^{\infty} F_Y(x) p_X(x) dx, \tag{9}$$

where $F_Y$ is the cumulative distribution function of Y. Then if we have samples, $x_n$, distributed according to $p_X(x)$ (i.e. $X \sim p_X(x)$), then

$$\hat{P} = \frac{1}{N_{\text{embed}}} \sum_{n=1}^{N_{\text{embed}}} F_Y(x_n) = \frac{1}{N_{\text{embed}}} \sum_{n=1}^{N_{\text{embed}}} p_{i,n} \tag{10}$$

is an estimate of this likelihood.

### Supplementary Figures

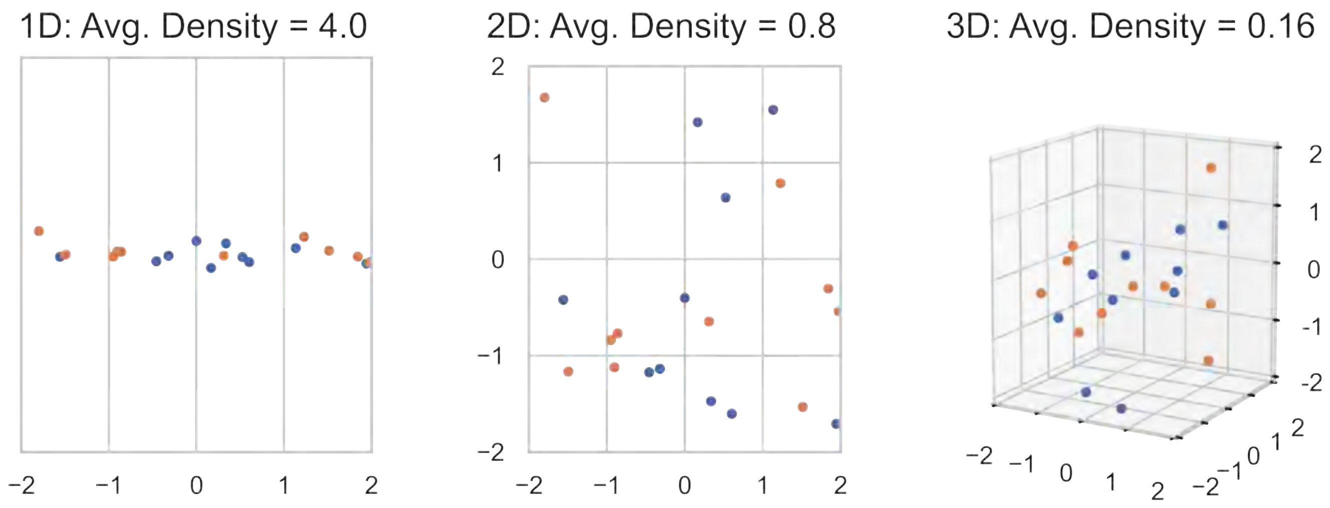


Figure S1: **The Curse of Dimensionality: Adding Axes Reduces Data Density** An illustration of the transition from 1D to 3D for simple random data. In each panel, the axes are divided into equally sized regions. As dimensions are added, the average density of samples in each region decreases from 20/5 = 4 data/unit to 20/25 = 0.8 data/unit to 20/125 = 0.16 data/unit. This exponential decrease in sample density is known as the “Curse of Dimensionality” [14] There are two colors of samples to illustrate what it might look like to differentiate two different hypothetical conditions.

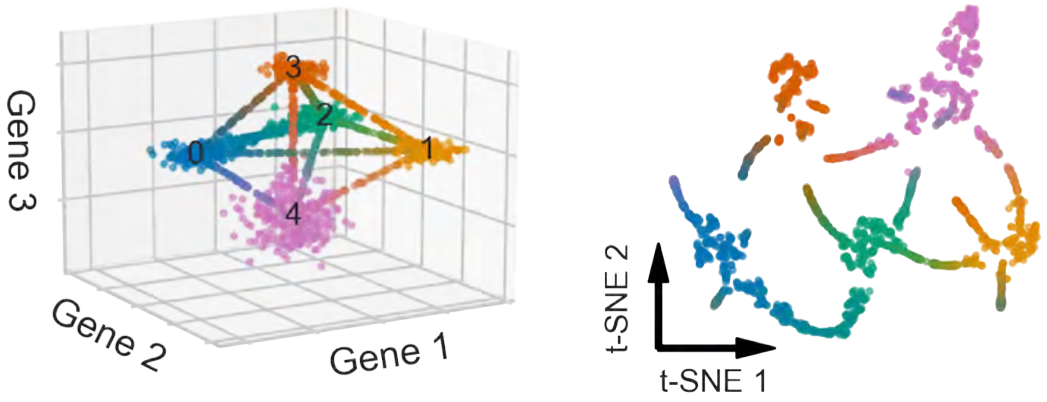


Figure S2: **Reducing the Dimensionality of a Tetrahedron Causes Distortions** (LEFT) Five clusters of "cells" of different sizes and shapes are centered on the vertices of a double tetrahedron . These clusters are also connected by small strands of cells. (RIGHT) The data are projected into 2D using t-SNE at default parameters. The pink and blue clusters, which were originally next to each other, have now been sent to opposite sides of the space. Similarly, the orange and yellow clusters have been artificially separated. The necessary symmetry breaking results in only cluster 2 (green) being connected to all other clusters in the lower-dimensional space.

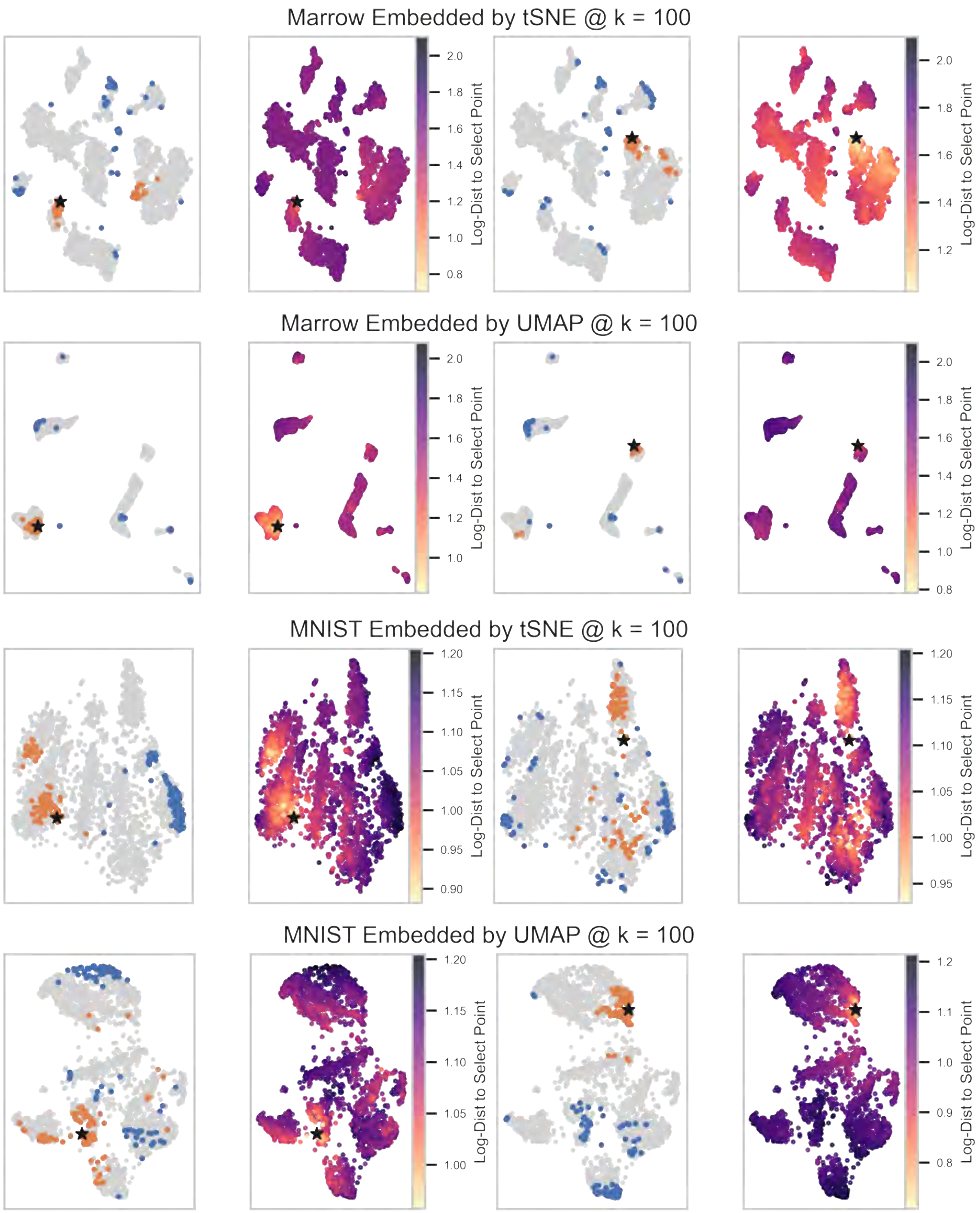


Figure S3: **Distances are not Uniformly Preserved by t-SNE or UMAP** The Tabula Muris Marrow [8] and MNIST digits [93] are embedded by both t-SNE and UMAP at $k_{\text{Eff}} \approx 100$ and $k = 100$, respectively. For each embedding of each data set, two random reference samples are selected (left two columns vs right two columns), shown as black stars. In the first and third columns, the location of the $k = 100$ nearest neighbors to the references in the original data are shown in the embedding in blue, and the location of the 100 furthest samples are shown in orange. The second and fourth columns show all samples colored by their log-distance to the reference in the high-dimensional data.

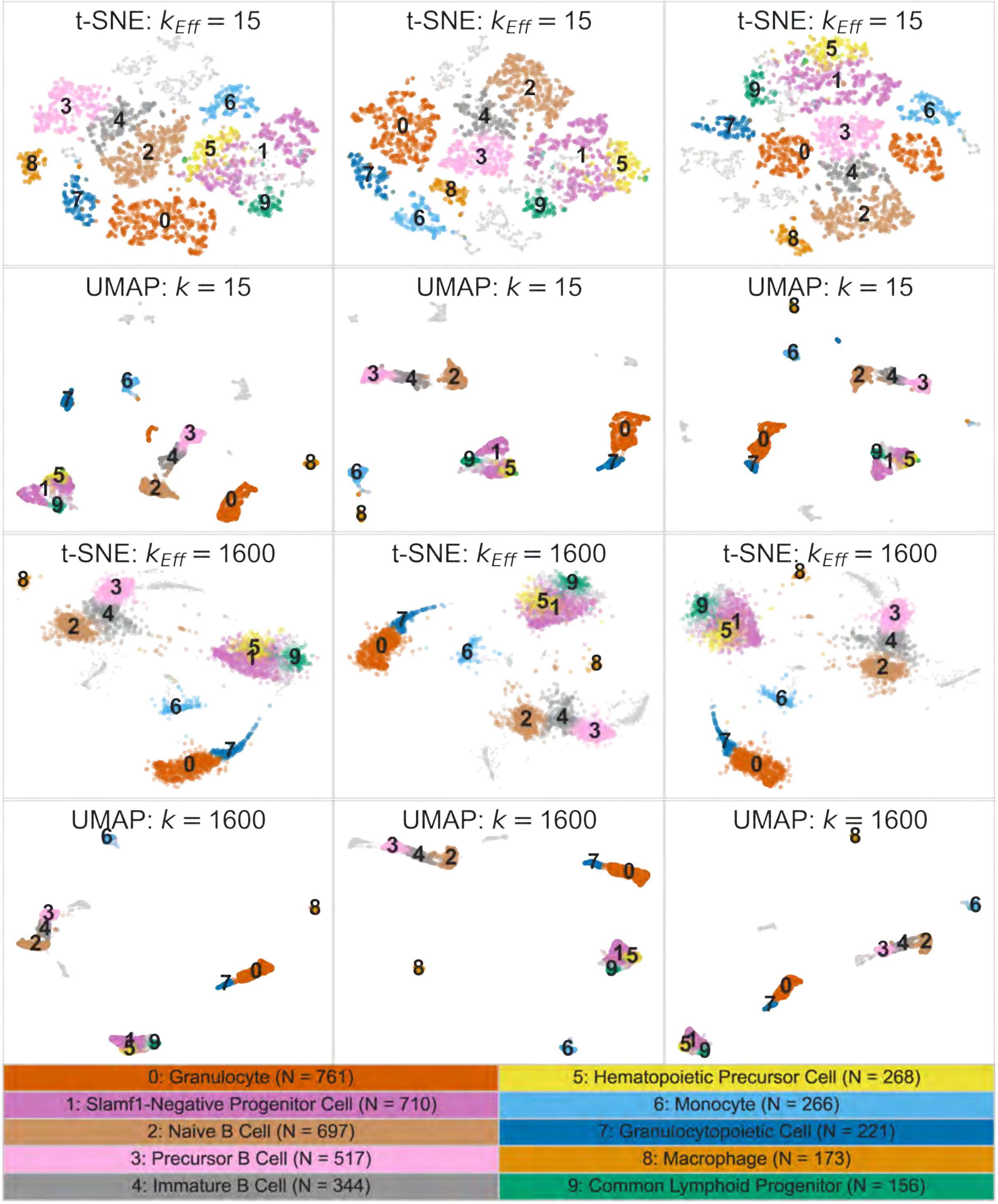


Figure S4: **Embeddings Generated by Stochastic Algorithms Change with Each Run (1):** Each of the parameter sets from Figure 1 generated with different random seeds. The coloring of the cells is the same as in Figure 1.

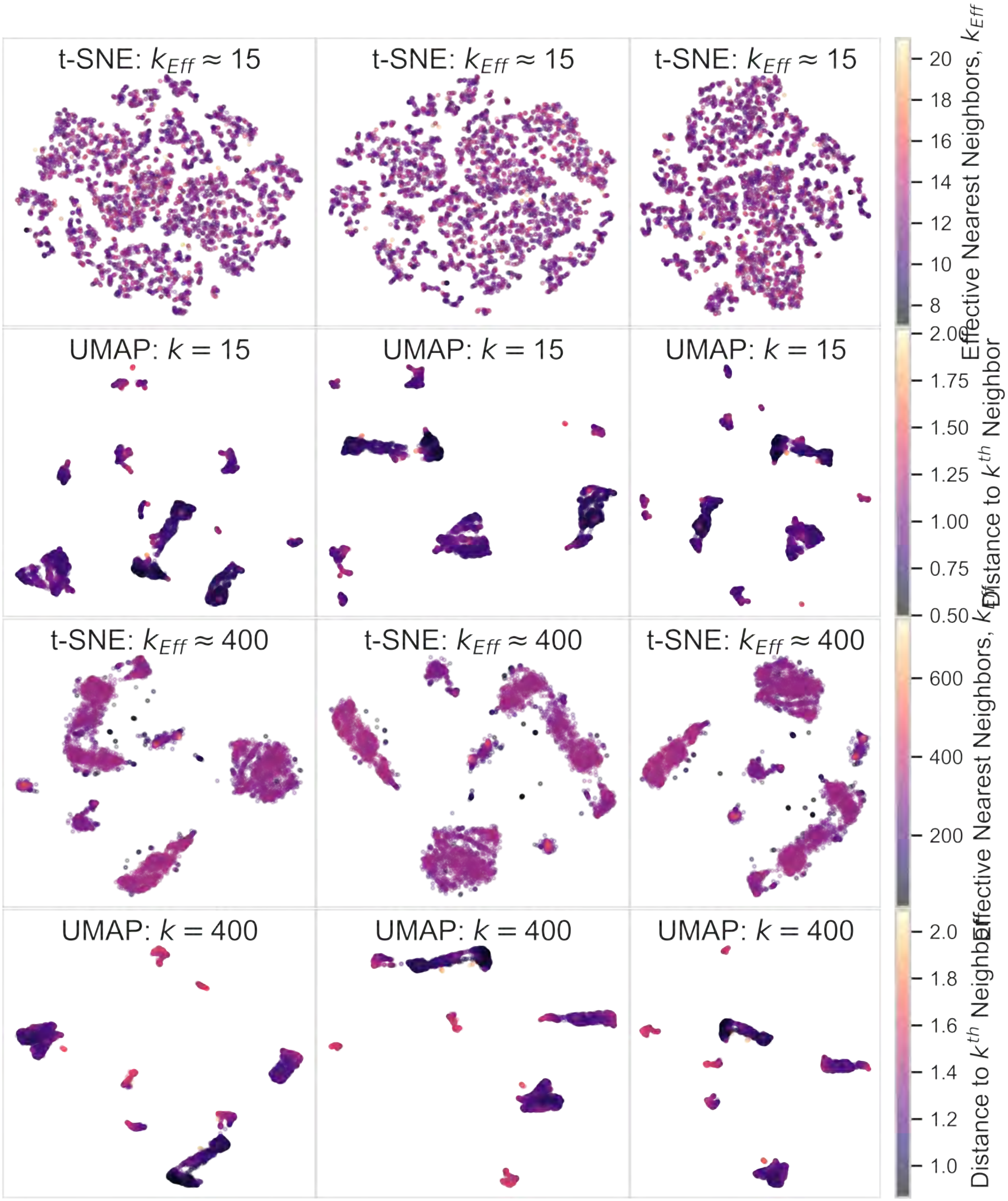


Figure S5: **The Structure of Data Varies Heterogeneously Across Embeddings:** Each of the parameter sets from Figure 1 generated with different random seeds. The cells in the t-SNE embeddings are colored by $k_{\text{Eff}}$ (see S3) and the cells in the UMAP embeddings are colored by the distance in the original data space to the $k$-th nearest neighbor.

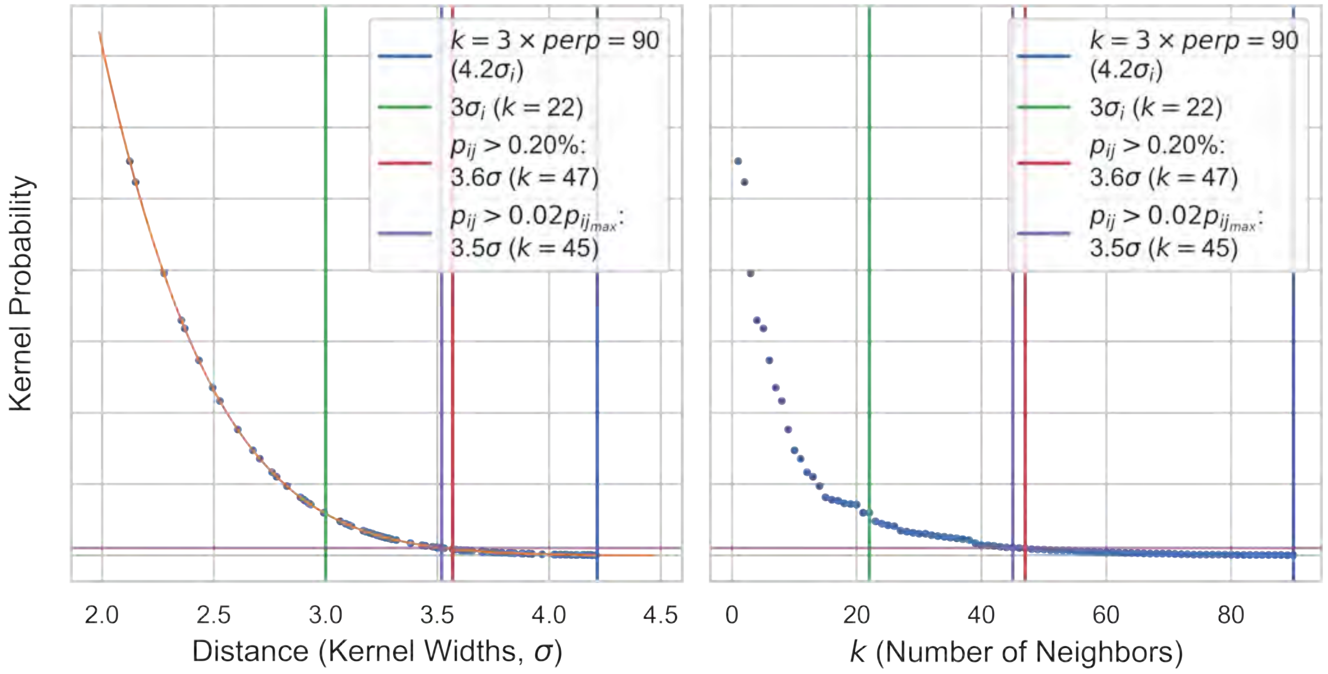


Figure S6: **Setting Thresholds on t-SNE's Kernel:** Using the method in [22] for setting the entropy of a Gaussian affinity kernel (`perplexity` = 30), the $3 \times 30 = 90$ nearest neighbors to an example cell can be visualized as a function of distance. Several putative thresholds can be set as proxies for "significantly contributing" neighbors: selecting a certain number of kernel widths, $\sigma_i$, shown in green; using a global threshold on the affinity values, shown in red; or using a local threshold based on the affinity of the nearest neighbor, shown in purple. In Section S3 we use the local threshold (purple) as our heuristic for assigning an "effective" number of nearest neighbors value to each value of `perplexity`.

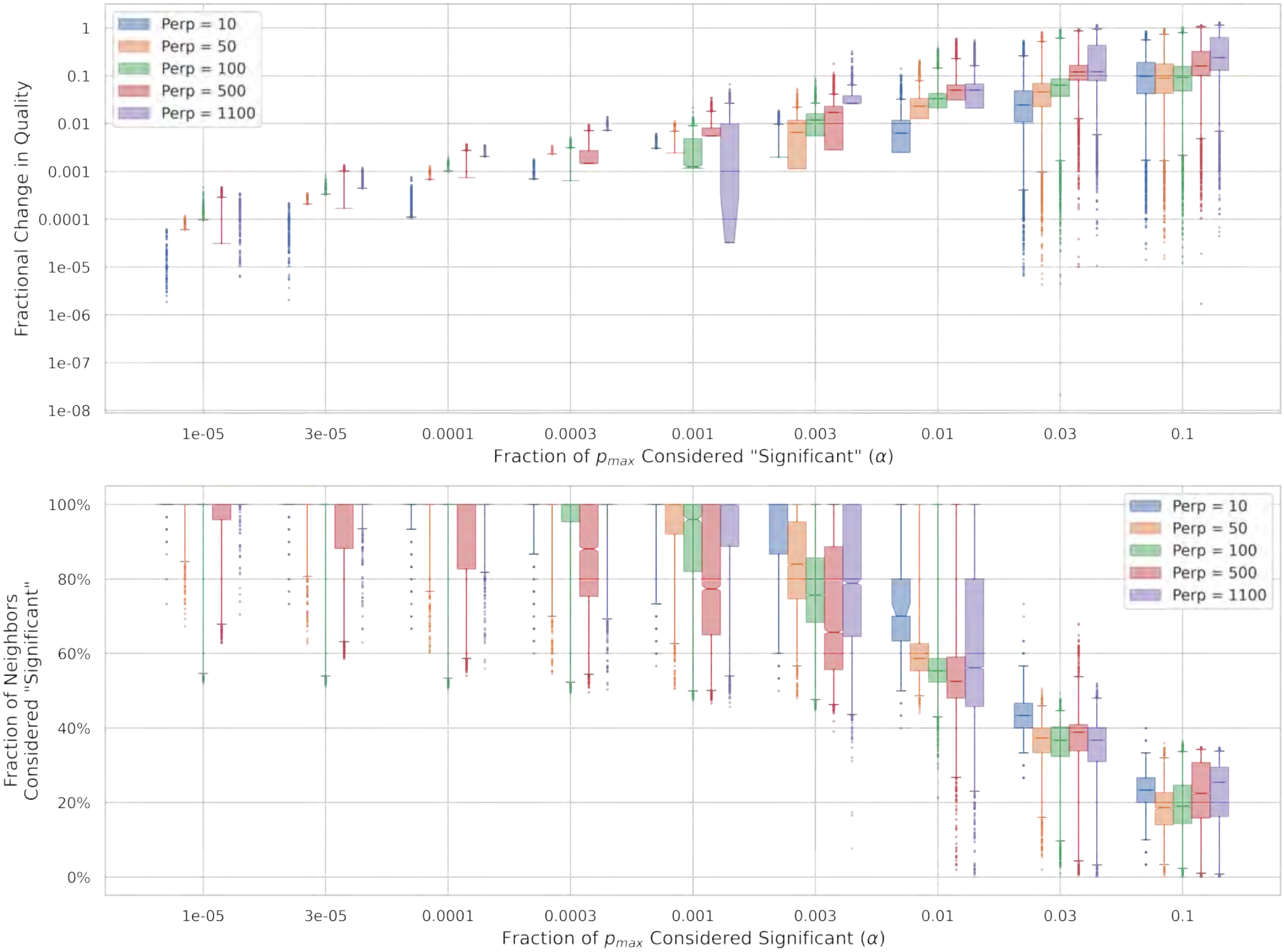


Figure S7: **Choosing an Appropriate Threshold for $k_{\text{Eff}}$:** (Top) The fractional change in $EES_i$ as a function of the local threshold, $\alpha$ for five different perplexities. (Bottom) The percentage of available (3×`perplexity`) neighbors that are considered "significant" at this threshold.

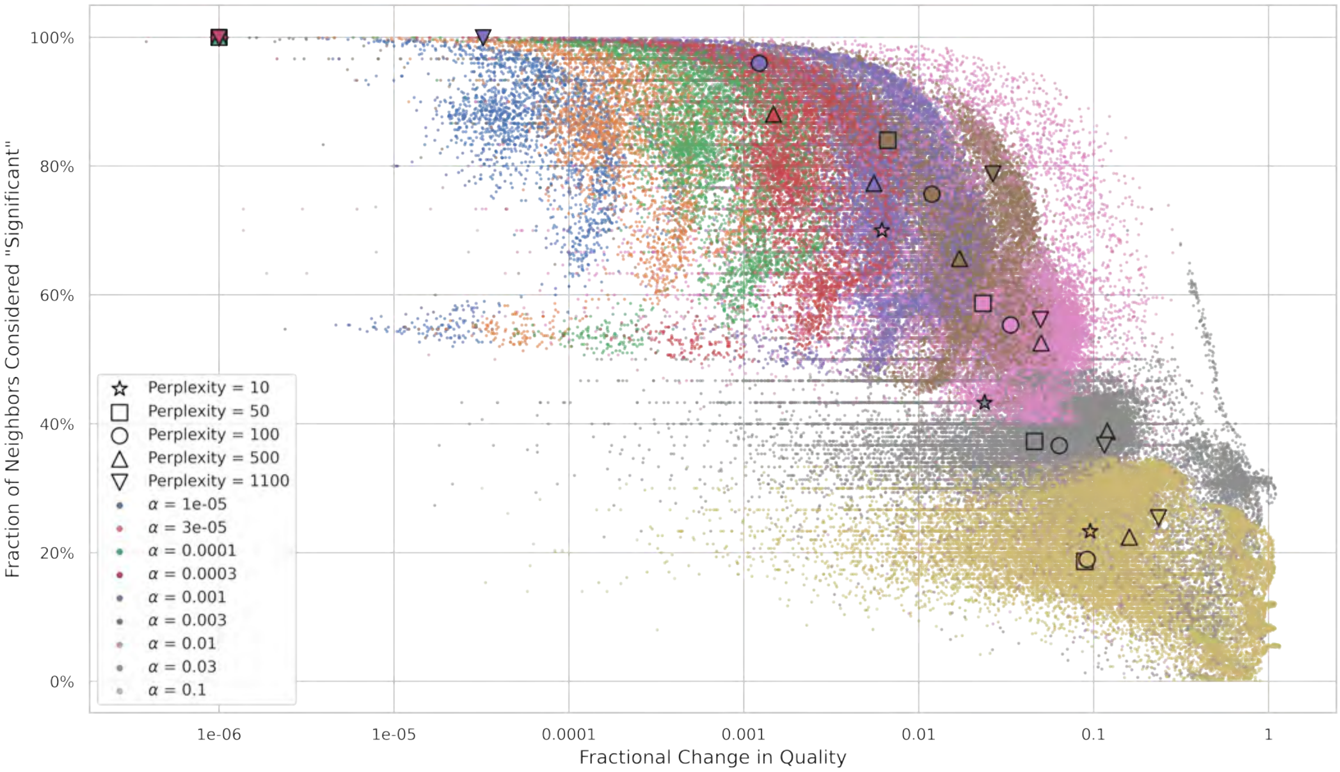


Figure S8: **Choosing an Appropriate Threshold for $k_{\mathrm{Eff}}$:** Directly plotting the fractional neighborhood consumption against the fractional change in $EES_i$ shows the expected pattern where as the threshold, $\alpha$, becomes more stringent (increases), fewer neighbors are considered significant, but there are greater changes to quality. The markers with black edges correspond to the median of that perplexity and threshold combination.

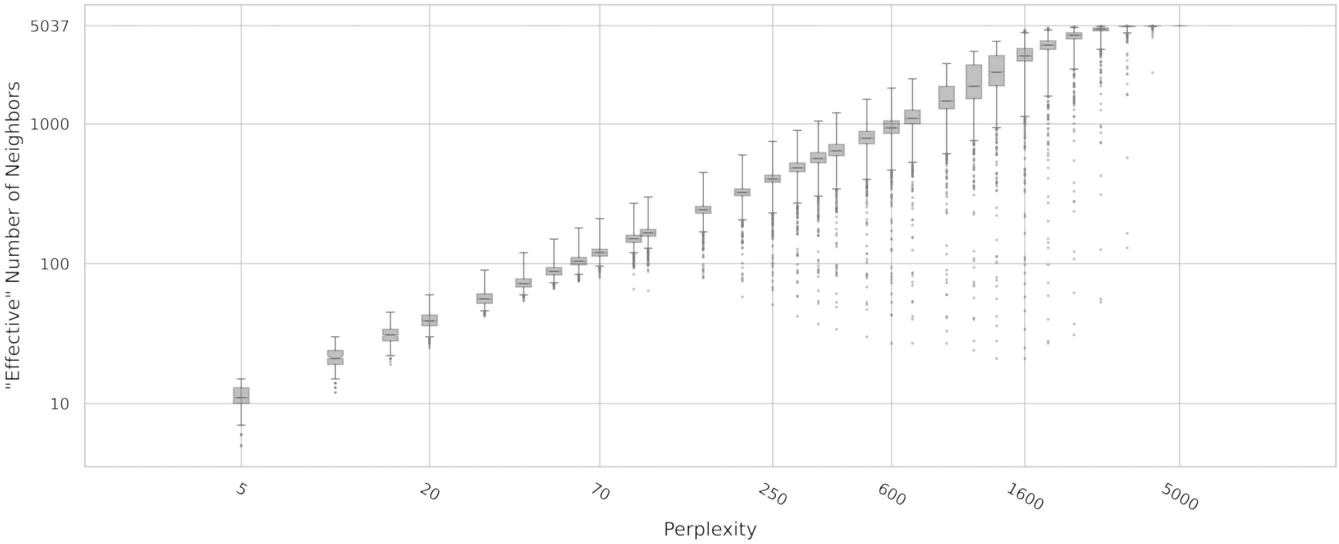


Figure S9: $k_{\mathrm{Eff}}$ **is Monotonically Related to Perplexity:** Once a threshold has been selected (here we choose $\alpha = 0.01$), the effective number of nearest neighbors, $k_{\mathrm{Eff}}$, can be calculated for each cell at each `perplexity`. Throughout the paper, we use the *median* $k_{\mathrm{Eff}}$ across cells to indicate a specific `perplexity`, since the relationship between $k_{\mathrm{Eff}}$ and `perplexity` is monotonic.

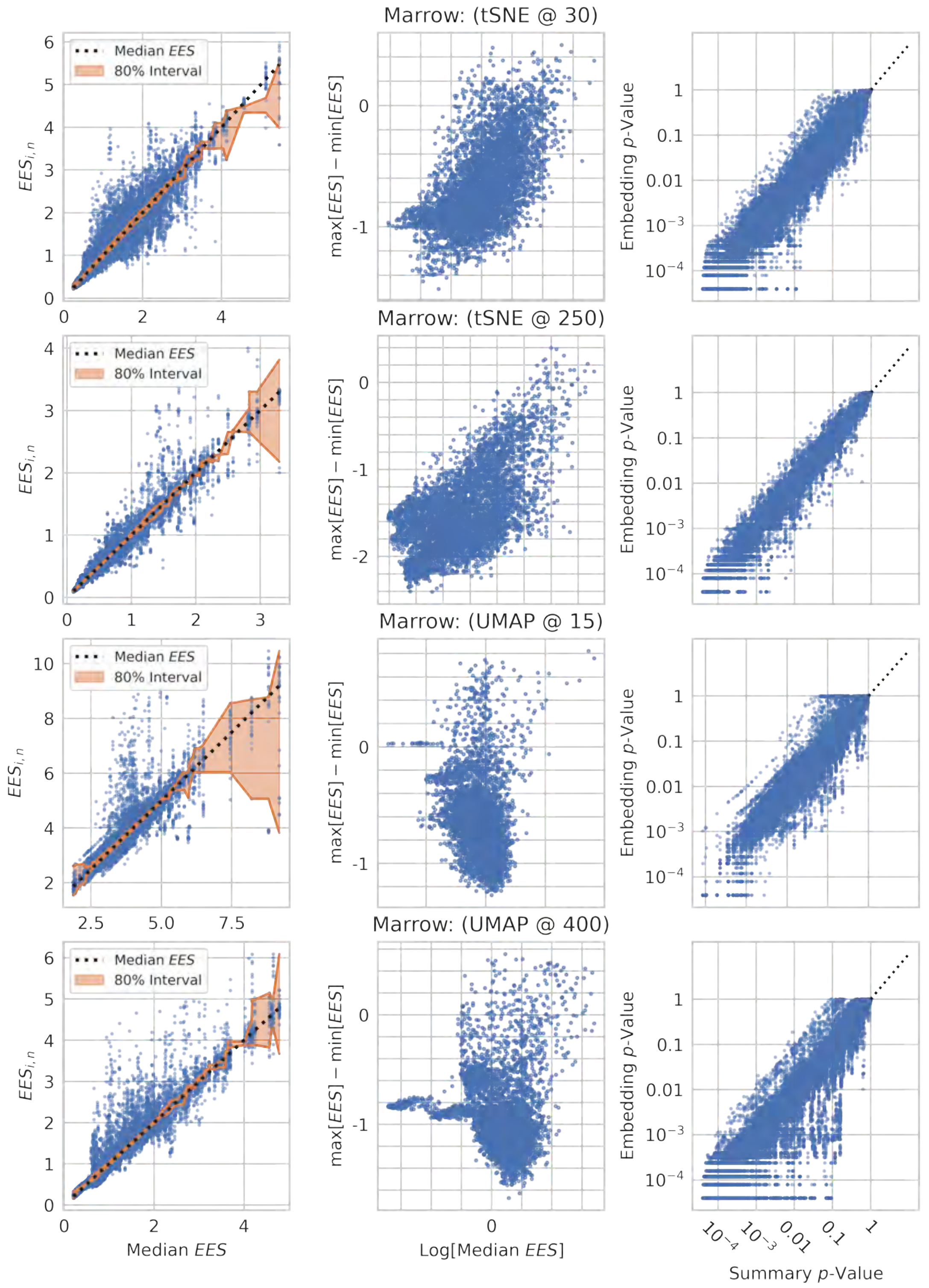


Figure S10: **Embedding Quality Varies Between Embeddings:** The Tabula Muris marrow data [8] were embedded 25 times by t-SNE and UMAP each at two hyperparameter values. The first column indicates the embedding quality of each sample in each of the 25 embeddings vs the median $EES_{i,n}$ for each cell across the embeddings. The rolling interval containing 80% of the cells is indicated in orange. The second column shows the range in a cell's embedding quality as a function of its median $EES_{i,n}$. The third column illustrates the relationship between the $p$-values calculated for each embedding and the consensus $p$-value found by averaging across embeddings.

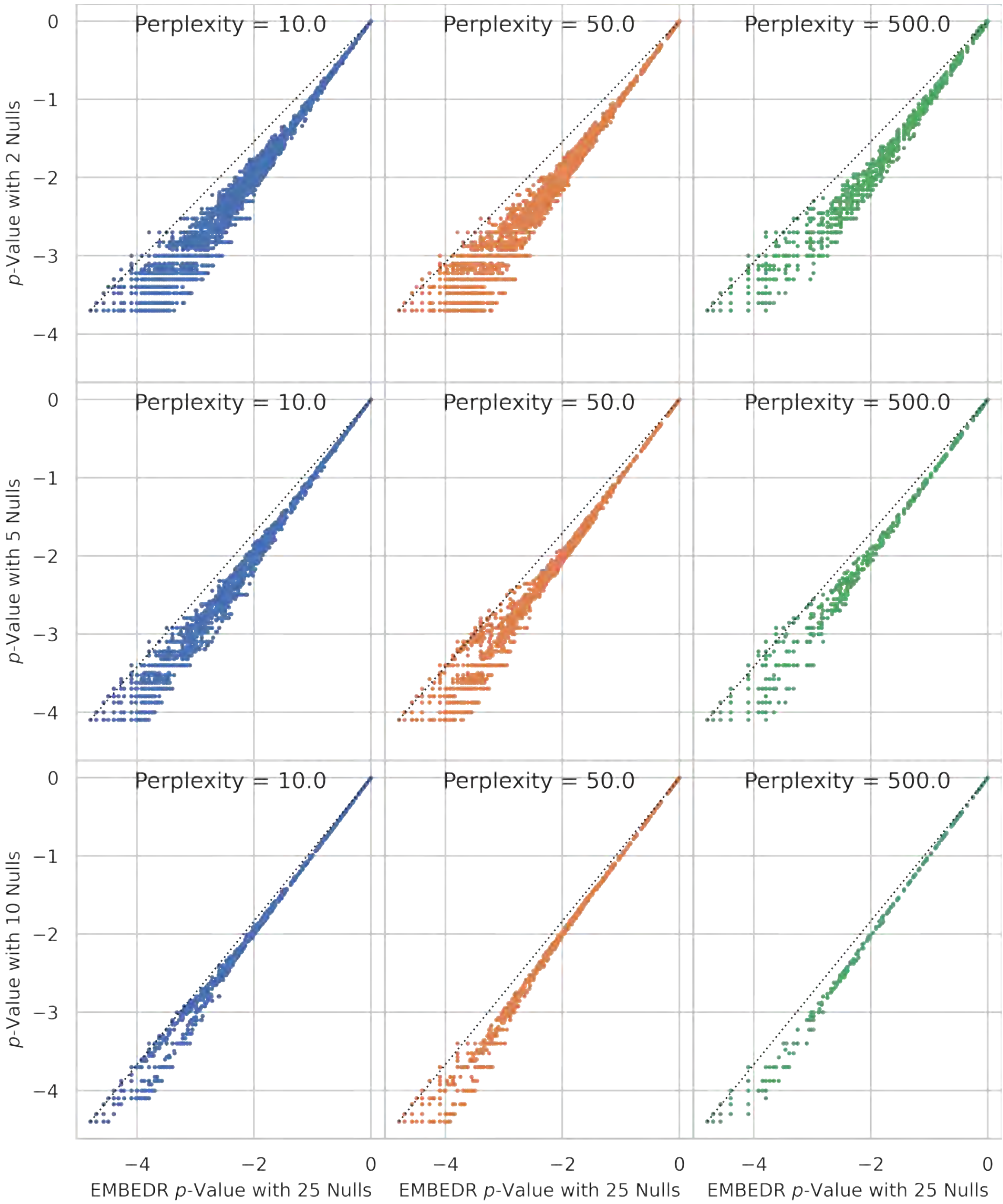


Figure S11: **Determining the Number of Resampled Nulls to Embed:** 25 Null data sets generated by marginally resampling the Tabula Muris marrow data [8] were embedded with t-SNE at several values of the t-SNE `perplexity` parameter. A consensus $p$-value was calculated by averaging as in Section S4 across all 25 embeddings and across sets of 2, 5, and 10 embeddings. These are compared in the top, middle, and bottom rows, respectively.

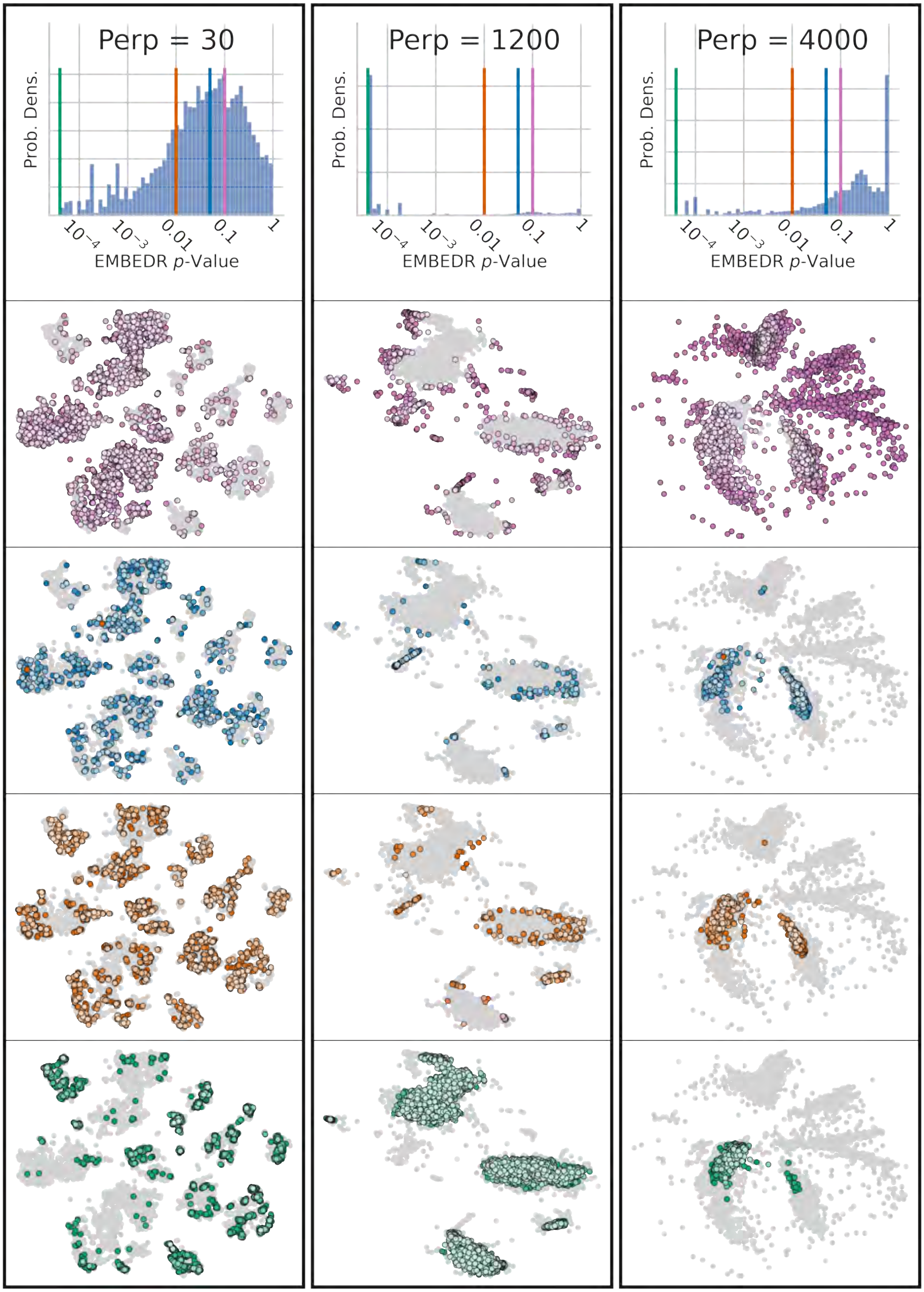


Figure S12: **Examining Cells at Different $p$-Value Levels:** The embeddings from Figure 4 are plotted, but only cells within each qualitative region of the $p$-value colorbar are shaded. (Top Row) The density of $p$-values alongside the 4 qualitative cutoffs are shown.

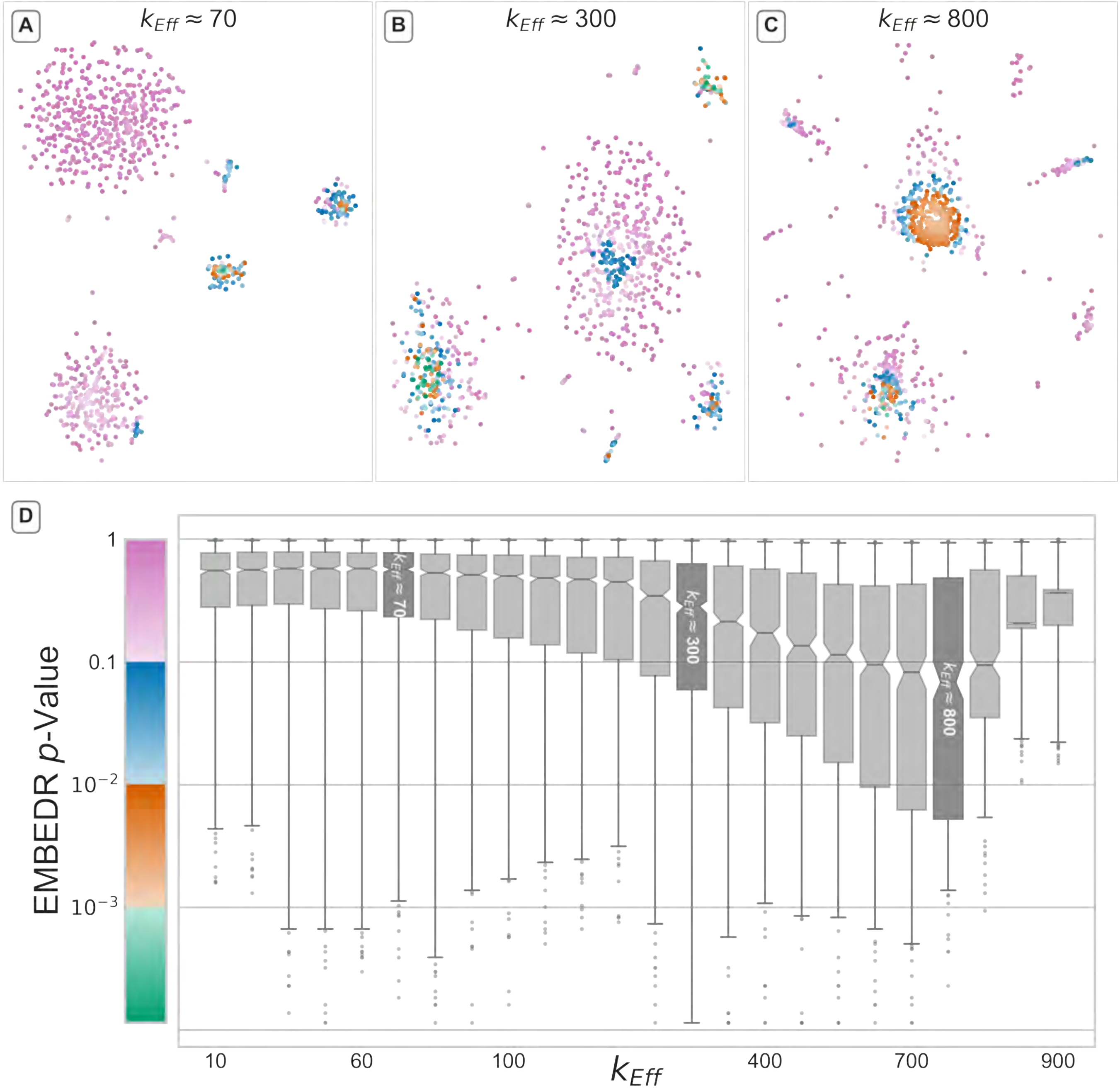


Figure S13: **A Global Sweep of $k_{\mathrm{Eff}}$ for Tabula Muris Diaphragm Tissue:** Another tissue from the Tabula Muris consortium from the diaphragms of several mice were embedded using t-SNE at a variety of scales as in Figure 4. The global minimum in EMBEDR $p$-value occurs at $k_{\mathrm{Eff}} \approx 800$ (`perplexity` = 600).

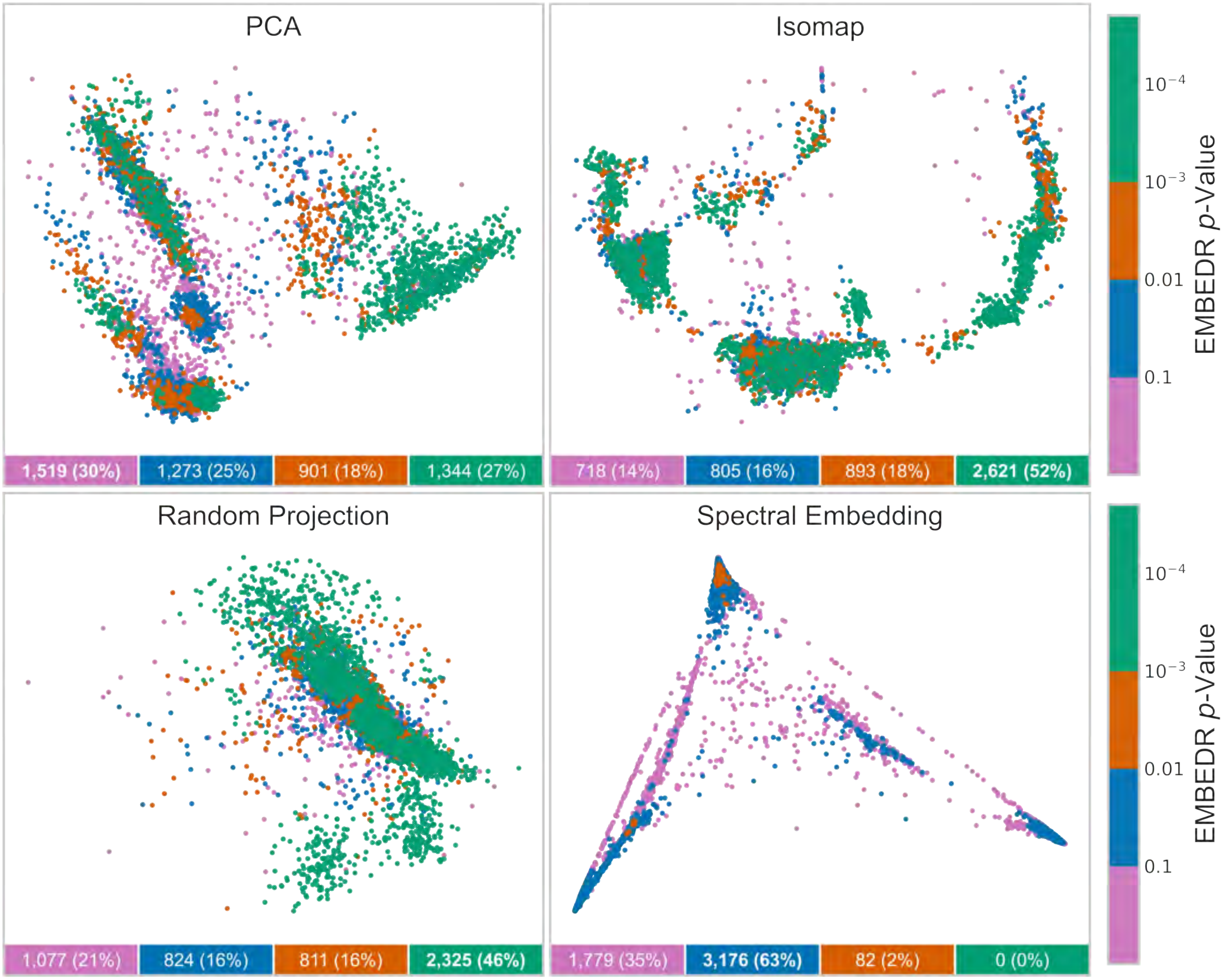


Figure S14: **EMBEDR Can be Applied to Arbitrary DR Methods:** EMBEDR is applied to PCA, Isomap [26], Sparse Random Projections [94], and Laplacian Eigenmaps [28] as they embed the Tabula Muris marrow data. The data were embedded once by each of these methods and the $p$-values were generated using 10 embeddings of null data. The scale of the affinity matrices used to calculate EES is set to correspond to $k_{\text{Eff}} \approx 400$, as this corresponds to a natural scale (Figure 4).

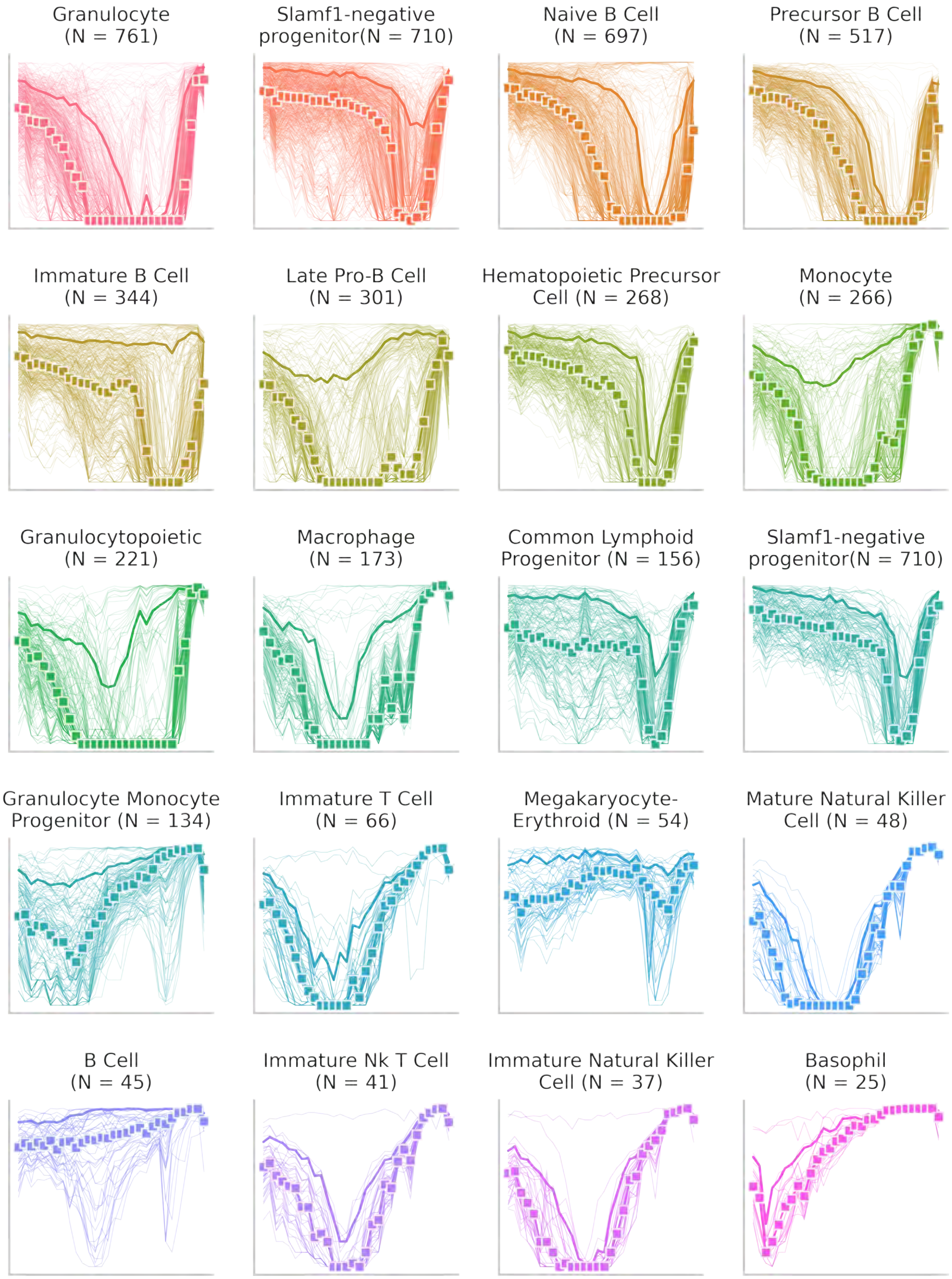


Figure S15: **A Sweep of $k_{\text{Eff}}$ for each Marrow Cell Annotation:** The 24 annotated cells types with the most cells from the Tabula Muris data set are embedded at several scales as in Figure 4. The spectra for each annotation is shown here, revealing a diversity of behaviors.

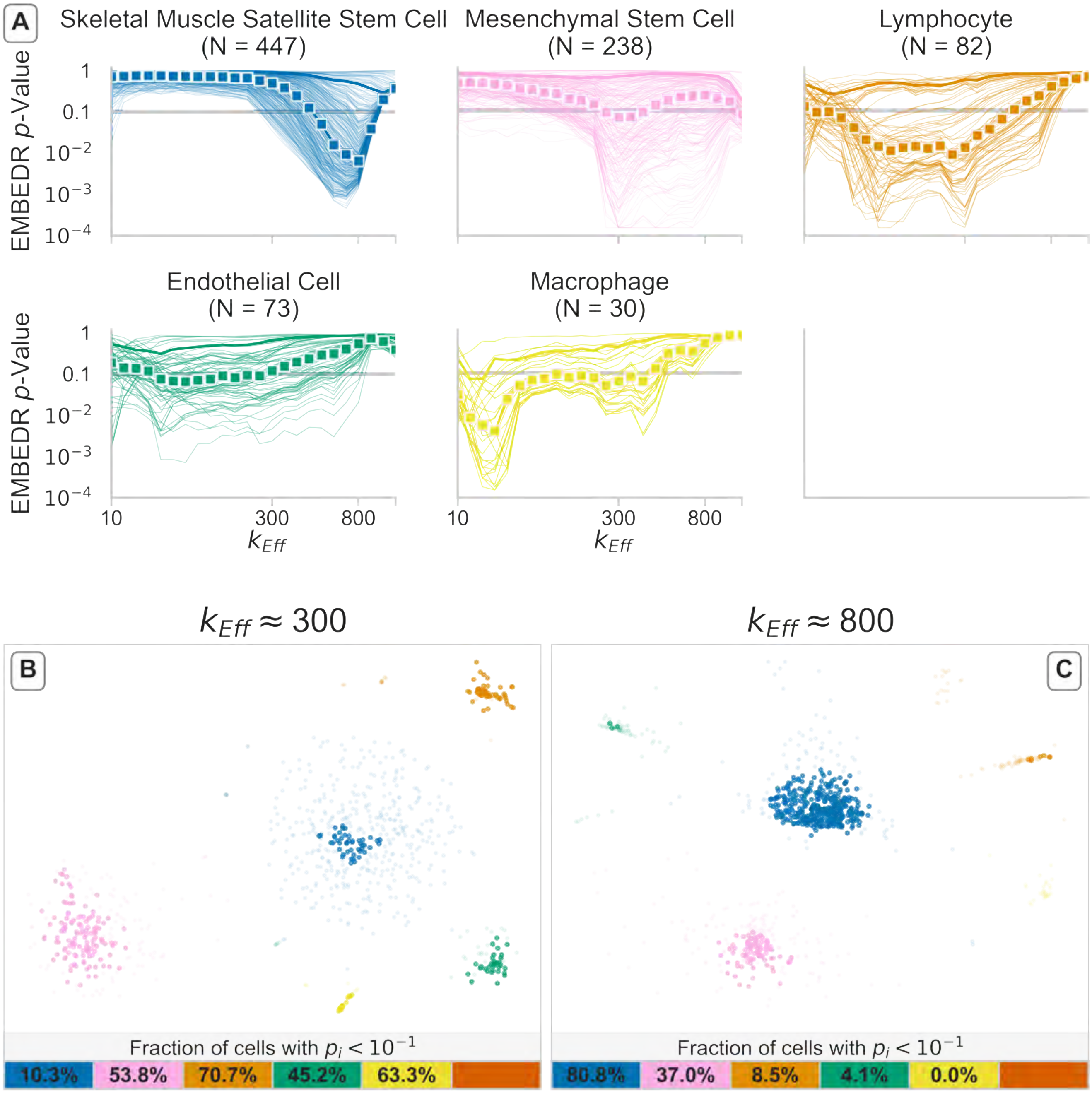


Figure S16: **A Sweep of $k_{\text{Eff}}$ for each Diaphragm Cell Annotation:** Figure 6 is repeated using the Tabula Muris diaphragm data. Here it can be observed that each annotated cell type is best embedded at different scales.

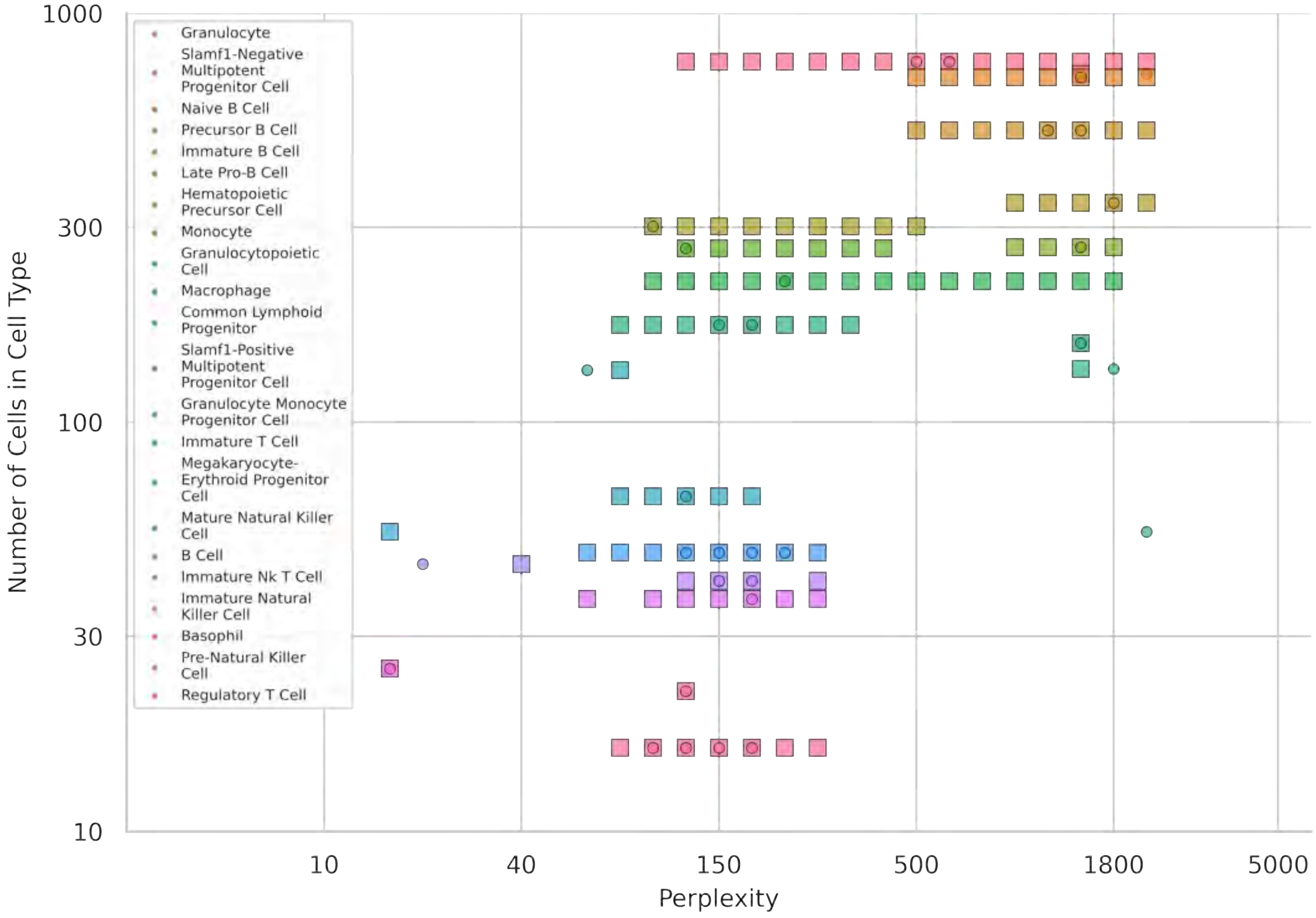


Figure S17: **Number of Cells in an Annotation Correlates with the Optimal $k_{\text{Eff}}$:** As a verification that cluster size is correlated with optimal embedding scale, the number of cells in a cell type annotation are compared to the location of the minimal median $p$-value for that annotation. These are shown as colored boxes. When the median was minimal for multiple values of `perplexity`, all such values of `perplexity` are indicated. The location of where the 90$^{\text{th}}$ percentile $p$-value was found is indicated by a colored circle. Since `perplexity` monotonically corresponds to effective neighborhood size, $k_{\text{Eff}}$, (see Figure S9) the desired relationship corresponds to larger cell annotations having larger optimal `perplexity` values, as seen here. This can also be observed in Figure S16, where the location of the minimal $p$-values increase with the larger cell annotations.
